# Distinct inflammatory and transcriptomic profiles in dominant versus subordinate males in mouse social hierarchies

**DOI:** 10.1101/2021.09.04.458987

**Authors:** Won Lee, Tyler M. Milewski, Madeleine F. Dwortz, Rebecca L. Young, Andrew D. Gaudet, Laura K. Fonken, Frances A. Champagne, James P. Curley

**Author notes:** Corresponding author: Dr. James P. Curley, Department of Psychology University of Texas at Austin Austin, USA.

## Abstract

Social status is a critical factor determining health outcomes in human and nonhuman social species. In social hierarchies with reproductive skew, individuals compete to monopolize resources and increase mating opportunities. This can come at a significant energetic cost leading to trade-offs between different physiological systems. Particularly, changes in energetic investment in the immune system can have significant short and long-term effects on fitness and health. We have previously found that dominant alpha male mice living in social hierarchies have increased metabolic demands related to territorial defense. In this study, we tested the hypothesis that high-ranking male mice favor energetically inexpensive adaptive immunity, while subordinate mice show higher investment in innate immunity. We housed 12 groups of 10 outbred CD-1 male mice in a social housing system. All formed linear social hierarchies and subordinate mice had higher concentrations of plasma corticosterone (CORT) than alpha males. This difference was heightened in highly despotic hierarchies. Using flow cytometry, we found that dominant status was associated with a significant shift in immunophenotypes towards favoring adaptive versus innate immunity. Using Tag-Seq to profile hepatic and splenic transcriptomes of alpha and subordinate males, we identified genes that regulate metabolic and immune defense pathways that are associated with status and/or CORT concentration. In the liver, dominant animals showed an up-regulation of specific genes involved in major urinary production and catabolic processes, whereas subordinate animals showed an up-regulation of genes promoting biosynthetic processes, wound healing, and proinflammatory responses. In spleen, subordinate mice showed up-regulation of genes facilitating oxidative phosphorylation and DNA repair and CORT was negatively associated with genes involved in lymphocyte proliferation and activation. Together, our findings suggest that dominant and subordinate animals adaptively shift energy investment in immune functioning and gene expression to match their contextual energetic demands.

**Highlights:**

- Immunity is shaped by stress and energetic pressures associated with social status
- Dominant and subordinate mice favor adaptive and innate immunity, respectively
- Dominants increase expression of genes involved in energy production
- Wound healing and DNA repair genes are upregulated in subordinates
- Genes related to maintaining and signaling social status are upregulated in dominants

## 1. Introduction

Social status is a significant factor affecting health outcomes in human and non-human social animals (Goymann and Wingfield, 2004; Sapolsky, 2004). One way social status impacts health is by shaping immune function (Bartolomucci et al., 2001; Habig and Archie, 2015; Lea et al., 2018; Snyder-Mackler et al., 2016). For animals living in social dominance hierarchies, social status is linked to specific behavioral repertoires and access to energetic resources, territory, and mates. For example, higher-ranking males tend to monopolize access to food and consume more than lower-ranking males; however, they also tend to expend more energy patrolling territory and investing in reproductive behavior than lower-ranking males (Lee et al., 2018, 2017; O’Connell and Hofmann, 2011). Disparate access to resources and energy expenditure associated with social status impacts the amount of energy available to maintain the immune system (Klein and Nelson, 1999). In this way, social status shapes how finite energy is allocated amongst the immune system and other sub-physiological systems such as reproduction, locomotion, thermogenesis, and cellular maintenance.

The direction and degree to which an animal’s social status influences energy allocation to immune function and health outcomes varies by sex, species, and the specific social dynamics of their social group (Beery et al., 2020; Habig et al., 2018; Habig and Archie, 2015; Snyder-Mackler et al., 2020). In humans, low socioeconomic status (SES) is associated with an exaggerated inflammatory response to stress (Jiang et al., 2020), higher risk of infection and disease (Adler et al., 1994; Cohen et al., 2008; Forastiere et al., 2007), and mortality (Chetty et al., 2016; Elo, 2009). Similarly, low status female rhesus monkeys exhibit dysregulated hypothalamic-pituitary adrenal axis (HPA) activity (Tung et al., 2012) as well as an immune system biased towards a proinflammatory, antibacterial response and away from an antiviral response, the reverse of what is observed in high status females (Snyder-Mackler et al., 2016). In contrast, high status male baboons show a heightened inflammatory response to bacterial infection (Lea et al., 2018). Such species and sex-specific differences in the association between social status and immune phenotype suggests that social context is an important determinant of how energy is allocated towards immune function. The mechanisms by which social context influences immune function is yet to be precisely determined.

One proposed framework for how high social status leads to compromised immune function is the life-history trade-off hypothesis. Within this framework, high-status individuals direct energy away from immune processes and prioritize the energy towards immediate reproductive success, which can result in compromised long-term immune function and health (Mills et al., 2010, 2009; Sheldon and Verhulst, 1996). For example, male jungle fowl that invest heavily in sexual signals to attract females show lower levels of circulating lymphocytes (Zuk et al., 1995). In male bank voles, high reproductive effort is associated with increased parasite infection rates and low survival rates (Mills et al., 2010). Another explicative framework is the stress response hypothesis that proposes that low social status has a detrimental impact on immune function (Dhabhar, 2009; Glaser and Kiecolt-Glaser, 2005). When living in groups, low-rank individuals often demonstrate compromised immune systems and worse health outcomes, such as high mortality rates and slow wound healing, even without physical injury or with access to ample nutritional resources (Archie et al., 2012; Blanchard et al., 1985; Koolhaas et al., 1997). These outcomes are likely induced through low-rank individuals experiencing chronic psychosocial stress characterized by a loss of control, a lack of predictability over their social environment, and a loss of outlets for frustration (Sapolsky, 1994). The health outcomes of this chronic psychosocial stress have been demonstrated in the resident-intruder paradigm and the social defeat paradigm; socially defeated animals exhibit prolonged elevation of proinflammatory cytokines, diminished splenocyte proliferation, and slow wound healing (Archie et al., 2012; Bartolomucci et al., 2001; Engler et al., 2004; Hodes et al., 2014; McKim et al., 2016; Sapolsky, 2005). Notably, these effects are only observed in animals who are susceptible to social defeat stress, as characterized by social avoidance following the contest, and not in resilient animals (Hodes et al., 2014; McKim et al., 2016). These hypotheses are not necessarily mutually exclusive as high and low status may both impact immune functioning negatively depending on the current social environment.

In this study, our aims were to investigate whether the daily energetic demands and stressors associated with social status shape immune function. We used an established social dominance paradigm in which cohorts of 10-12 male CD-1 mice live in an ethologically relevant group housing system and reliably form stable linear hierarchies (So et al., 2015; Williamson et al., 2016). As a social hierarchy is established, each animal occupies a unique social rank and dramatically different behavioral phenotypes emerge between ranks. The highest ranking male (alpha) exhibits higher rates of aggression compared to all other individuals, is more active in patrolling territory and eats and drinks more frequently (Lee et al., 2018). Alpha males also face further metabolic energy demands as they increase their production of major urinary proteins (MUPs) in the liver, which are then deposited for scent-marks and used to signal their dominance status to other animals (Lee et al., 2017; Nelson et al., 2015). Ranked below the alpha are the sub-dominant individuals who initiate aggression towards relatively subordinate individuals despite repeatedly receiving high levels of aggression from relatively more dominant individuals. Subordinates are the lowest ranking individuals who rarely show aggression. While these status-related phenotypes are typically consistent across cohorts, there can be some variation in the degree of aggressiveness (i.e., despotism) exhibited by alphas. Highly despotic alpha males exert frequent aggression and inhibit aggressive behavior of all other group members, and notably, subordinate individuals living with highly despotic alpha males exhibit elevated basal plasma corticosterone (CORT) concentrations (Williamson et al., 2017).

This accumulating evidence demonstrates the social dominance paradigm is uniquely suited to investigate the relationship between social status, metabolism, and immune function. We hypothesized that the energetic demands of establishing and maintaining dominance in a hierarchy would reshape immune profiles such that energy allocation would be polarized towards cheaper adaptive immune components and less towards more energetically expensive innate immune components (Habig et al., 2018; Habig and Archie, 2015; Lee, 2006). In accordance with this hypothesis, we predicted the occurrence of shifts in metabolism markers indicative of higher energy expenditure. Moreover, we predicted this same framework would hold true for more subordinate individuals, such that they would have enough energy to allocate towards innate immune components as it would facilitate recovery from injuries (Archie et al., 2012; Kiecolt-Glaser et al., 1995). To test the hypotheses, we measured immune cell composition and plasma CORT from peripheral blood of male CD-1 mice before and after the formation of social hierarchies. As a proxy measurement of long-term stress exposure (Lee et al., 2010; Lee et al., 2011), we measured *Fkbp5* DNA methylation levels. Lastly, we profiled the hepatic and splenic transcriptomes to investigate the expression pattern of genes involved in energy metabolism and the immune system function. Taken together, this study adds multifaceted insights on the physiological mechanism of how social information is translated into adaptive energy allocation to sub-physiological systems essential to reproductive efforts and long term health.

## 2. Materials and Methods

### 2.1 Behavioral experiment procedure

The timeline and behavioral procedures are summarized in **Fig. 1A**.

**Figure 1.**
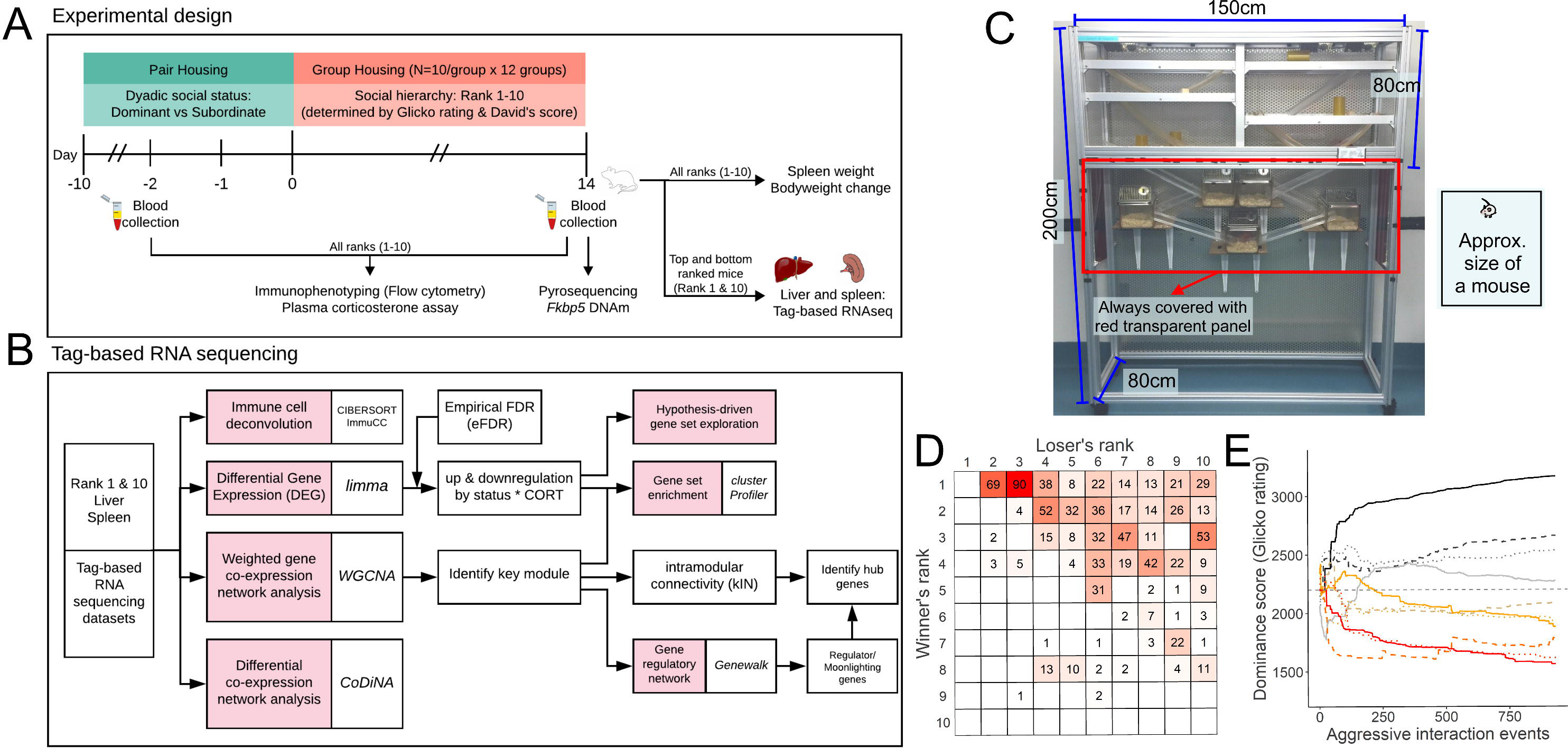
(A) Study design. Mice were housed in pairs for 10 days then assigned to a social group of 10 male mice in a vivarium (C) designed to mimic wild mouse habitat. We determined individual social ranks based on daily behavioral observations of aggressive interactions. On pair housing Day 9 and on Group housing day 14, blood samples were collected from all subjects to measure immunophenotypes and plasma corticosterone levels. Also at Group housing day 14, liver and spleen tissue samples were collected from Rank 1 (Alpha) and Rank 10 (the most subordinate) from each social group and processed for whole transcriptome data via (B) Tag-based RNA sequencing. A sociomatrix (D) and temporal dynamics (E) of individual Glicko ratings of one exemplar social group (Cohort I).

#### 2.1.1 Animals

A total of 120 male outbred CD-1 mice were used in the current study. Mice at 7 weeks of age were obtained from Charles River Laboratories (Wilmington, MA, USA) and housed in pairs for 10 days in standard sized cages (27 × 17 × 12 cm) surfaced with pine shaving bedding (Nepco, Warrensburg, NY, USA). Standard chow and water were provided ad libitum. All mice were individually marked with a blue non-toxic animal marker (Stoelting Col., Wood Dale, IL, USA). All experiments and housing mice were approved and conducted in accordance with the University of Texas at Austin Institutional Animal Care and Use Committee (IACUC Protocol No. AUP-2018-00119) and National Health Institute (NIH; Bethesda, MD, USA) animal care guidelines.

#### 2.1.2 Housing

Mice were housed in the animal facility in the Department of Psychology at the University of Texas at Austin on a 12:12 light-dark cycle with white light on at 2300h and red light on at 1100h. The room was kept under constant temperature (21–24 °C) and humidity (30–50%). Animals were housed in pairs for 10 days and their dyadic dominant-subordinate status were determined via observation (see below). At Zeitgeber time 11 (ZT11) on the 11^th^ day of pair housing (PD11 = group housing day GD01), groups of 10 mice were weighed and assigned into group housing vivaria (see **Fig. 1A**). We assigned mice from each pair to different social groups so that all animals had no previous experience with any other animal in the social group. Each social group consisted of 5 dominant males and 5 subordinate males from pairs. The group housing vivaria (**Fig. 1C**) were constructed as previously described (Williamson et al., 2016). Briefly, each vivarium consists of an upper level with multiple shelves (total available surface: 36,000 cm^2^ = 3 floor x 150 cm x 80cm) and a lower level with five connected nest boxes (2,295 cm^2^ = 5 cages ×27 cm ×17 cm). The total surface of a vivarium is 62,295 cm^2^ (6,230 cm^2^ per mouse) and was covered in pine shaving bedding. Food and water were provided at the top of the vivarium ad libitum and mice could access all levels of the vivarium through interconnected tubes, nest boxes and shelves.

#### 2.1.3 Submandibular blood collection

At ZT06 on day 9 of pair housing (PD09), 150-200ul of blood was collected from each mouse via submandibular puncture as described in Golde et al., (2005). Once enough blood was collected 2-3 seconds after the puncture, the puncture site was immediately pressed gently with sterile gauze to stop bleeding. Mice were monitored thoroughly for any sign or injury or pain after they returned to home cages. Blood samples were collected in EDTA-coated BD Microtainer tubes (Becton Dickinson, and Company, Franklin Lakes, NJ), gently inverted a few times then stored on ice for 30-60 minutes before subsequent procedures. 100ul of blood aliquots were used for flow cytometry. The rest of blood samples, usually ∼50ul, were centrifuged (14,000xg for 10 min at 4°C). Plasma was collected and stored at -80°C.

#### 2.1.4 Tissue harvest and trunk blood collection

At ZT06 on day 15 of group housing (GD15), mice were retrieved from the group housing system. Mice were weighed and euthanized via decapitation. Trunk blood samples were collected into EDTA-coated Vacutainer tubes (Becton Dickinson) and gently inverted a few times and stored on ice for 45-60 minutes prior to subsequent procedures. As above, 100ul aliquot of blood samples were used for flow cytometry. 200ul of blood aliquots were centrifuged to collect plasma (stored at -80°C until corticosterone assay), then subsequently processed for gDNA extraction within 4 hours of the collection. Anogenital distance, the distance between the center of the anus and the base of the genitalia, was measured with a digital caliper as previously described (Ophir and delBarco-Trillo, 2007). Spleen was dissected carefully to exclude fat mass attached to it and weighed with a highly sensitive scale with accuracy level to 0.0001g. Livers and spleens were fresh frozen in a hexane cooling bath chilled on dry ice then stored in -80°C until RNA extraction.

#### 2.1.5 Social behavior observations

All behavioral observations were conducted during the dark cycle as described previously (Lee et al., 2017; So et al., 2015; Williamson et al., 2016). Briefly, during pair housing, behavioral observations were undertaken 1 hour each day to determine dyadic dominant-subordinate status. After 7 days of observation, we determined a mouse in a pair to be dominant if a mouse had more than three consecutive wins over its cagemate. During the 14 days of group housing, an average of 2.12 hours of daily behavioral observations were conducted on each social group. Trained observers used all occurrence sampling to record social interactions between two mice. In detail, the observers recorded the initiator and the responder of each social interaction, and specific behavior(s) exhibited by each animal based on the ethogram presented in Supplemental Table S1. Data were collected using either Android devices or Apple iPad and directly uploaded to a timestamped Google Drive via a survey link.

### 2.2 Biological sample processing methods

#### 2.2.1 Genomic DNA extraction and bisulfite pyrosequencing

200ul aliquots of blood samples collected on GD15 were used for genomic DNA (gDNA) extraction for bisulfite pyrosequencing using MasterPure DNA Purification kits (Lucigen, Middleton, WI, USA) according to the manufacturer’s protocol. We used a Take3 plate and Biotek multimode-microplate reader to determine DNA concentration. 500ng of gDNA for each sample was submitted to EpigenDX (Hopkinton, MA, USA) for standard bisulfite pyrosequencing procedures with primer sets developed by EpigenDX. Based on previous studies (Ewald et al., 2014; Kitraki et al., 2015; Y. S. Lee et al., 2010), we chose the assay that measures DNA methylation levels across three CpGs site located ∼200bp upstream of Exon 5 of *Fkbp5* gene.

#### 2.2.2 Corticosterone enzyme immunoassays

Plasma samples were diluted 1:125 fold with assay buffer and were run in duplicate according to the manufacturer’s instruction (K-014, Arbor assays, Ann Arbor, MI).

#### 2.2.3 Immunophenotyping using flow cytometry

We designed two-panel multicolor flow cytometry analysis based on a peer-reviewed immunophenotyping protocol, Optimized Multicolor Immunofluorescence Panel (OMIP), to maintain the validity and reproducibility of immunophenotyping analysis. We adapted the staining protocol and gating strategy provided in OMIP-032, Unsworth et al. (2016) with a few modifications. 150ul aliquots of blood samples collected on PD09 or GD15 were lysed with 2ml of red blood cell lysis buffer (Qiagen, Hilden, Germany) in 5ml Eppendorf tubes for 5 minutes. Separated white blood cells were blocked with 1:100 Fc-Block (BD Biosciences) for 20 mins at 4°C in darkness. Each blocked sample was speared into two tubes and stained with cocktails of antibodies for 30 mins at 4°C in darkness. Two panels of antibody cocktails were designed to detect specific immune cells in the blood (Panel 1: neutrophil, macrophage, monocytes, lymphoid dendritic cells, myeloid dendritic cells; Panel 2: natural killer cells, B cells, Helper T cells, cytotoxic T cells). Detailed information on the antibodies used for fluorescent staining for specific cell markers are listed in Supplemental Table S2. Samples were acquired on BD LSR Fortessa X20 (BD Biosciences, East Rutherford, NJ, USA) maintained by the University of Texas at Austin Microscopy & Imaging Facility via FACS Diva software (BD Biosciences). We followed gating strategies described in Unsworth et al. (2016) which allows us to exclude dead cells and debris and to discriminate doublets from singlets. Raw data was analyzed via FlowJo v10 (TreeStar, Ashland, OR, USA).

#### 2.2.4 RNA extraction from liver and spleen tissue samples

For RNA sequencing, we selected livers and spleens of alpha (Rank 1) and the most subordinate mice (Rank 10) from each cohort based on their David’s score and Glicko rating (see below). The alpha male at GD15 from Cohort G was excluded from the analysis as they displaced the initial alpha male (see Fig. S2). We dissected ∼30mg of the liver and spleen into 2ml Reinforced Microvials (2007, BioSpec Products, Inc., Bartlesville, OK) containing 1mm zirconia/silicon beads (11079110z, BioSpec). Dissected tissue samples were homogenized in 500ul lysis buffer (Thermo Fisher Scientific, Waltham, MA; MagMax Total RNA isolation kit, Cat. No. A27828) with 0.7% beta-mercaptoethanol using the beads beater (607, Bio Spec). Lysates were incubated at room temperature for 5 minutes and 100ul aliquots of homogenates were proceeded to RNA extraction on the KingFisher Flex (Thermo Fisher Scientific, Cat. No. 5400630l) with an additional DNase step added according to the manufacturer’s protocol. RNA quality was determined using a RNA 6000 Nano Assay with BioAnalyzer (Agilent Technologies, Santa Clara, CA) and RNA concentration was determined with Quant-it RNA High Sensitivity assay kit (Thermo Fisher Scientific, Cat. No. Q33140). RNA samples were normalized to 100ng/ul and stored at -80°C before submission for sequencing. Extracted RNA samples were processed at the Genome Sequence and Analysis Facility at the University of Texas at Austin for Tag-based RNA sequencing. This method is a cost-effective and highly reliable approach specifically designed to measure abundances of polyadenylated transcripts yielding data for differential gene expression analysis in well annotated genomes (Lohman et al., 2016; Meyer et al., 2011). Libraries were constructed with a protocol described in Lohman et al. (2016). Reads were sequenced on the NovaSeq 6000 SR100 with minimum reads of 4 million and the target reads per sample of 5 million.

### 2.3 Statistical analysis

All statistical analyses were undertaken in R version 4.0.3 (R Core Team, 2021).

#### 2.3.1 Emergence of hierarchies and group dominance structure, and determination of individual dominance ranks, Glicko ratings, and David’s scores

For each social group, we calculated five dominance structure measures as previously described (Williamson et al., 2019, 2016). Briefly, we calculated Landau’s h value and triangle transitivity (ttri) to confirm the linearity of the social hierarchies. Both range from 0 to 1 with 1 indicating the hierarchy is perfectly linear. Significance tests were conducted and p-values determined using 10,000 Monte-Carlo randomizations (de Vries, 1995; McDonald and Shizuka, 2012) using the compete R package (Curley, 2016). We used despotism (Williamson et al., 2016) to determine the degree to which alpha males dominated all other individuals. Despotism is defined as the proportion of all wins by the alpha male of the group over the total number of aggressive interactions observed within the group. We used the Glicko rating system (Glickman, 1999) to confirm the emergence of individual ranks across observed aggressive interactions. Briefly, all individuals start with the same initial rating (2200). Then rating points are added for a winner or subtracted for a loser after each aggressive interaction. The degree of points added or subtracted is based on the difference in ratings between the winner and the loser (See Williamson et al., (2016) for more information). Finally, we measured the individual David’s score of individuals based on the win-loss sociomatrices. David’s score is a win proportion measure adjusted for opponent’s relative dominance in the hierarchy (Gammell et al., 2003) and was calculated using the compete R package (Curley, 2016). With David’s score, mice can be then further categorized into three distinct social status groups: alpha (rank 1, the highest David’s score), subdominant (David’s score >=0), and subordinate (David’s score <0) as previously described in Lee et al. (2017, 2018).

#### 2.3.2 Statistical analysis of biological measurements

All Bayesian (generalized) linear regressions were fitted using the R package brms (Bürkner, 2020). The convergence of each model was tested with the Gelman-Rubin convergence statistic and we confirmed all models converged (Rhat = 1). Throughout the manuscript, we discuss parameter estimates to be statistically significant when the 95% credibility interval (CI) does not overlap 0. Spleen weight is analyzed as relative weight to the body weight (absolute spleen weight/ body weight). Growth rate is defined as body weight change during group housing proportional to the original body weight ((body weight at GD14 - GD01)/GD01 *100). Proportion of each immune cell type was scaled and centered prior to analysis. We tested the association between the adjusted spleen weight or DNA methylation % of each CpG sites (CpG 1, CpG 2, CpG 3) against David’s score, frequency of wins or losses, growth rate with a Gaussian distribution with social group ID as a random factor. We tested if there is a difference in each cell proportion between dominant and subordinate mice within each pair from the blood collected at the end of the pair housing (PD09) with pair housing social status as a fixed factor and pair housing ID as a random factor. Similarly, we tested the effect of social dominance on immune cell proportion measured on GD14 with David’s score as a fixed factor and social group ID as a random factor. To see if the level of a specific type of immune cells in the peripheral blood prior to group housing (blood collected on PD09) predict the David’s score of each individual in the social hierarchies, we fitted proportion of each type of immune cells measured by flow cytometry against David’s score with a Gaussian distribution with social group ID and pair housing ID as random factors. We tested whether there is significant change in the proportion of cells from blood collected on PD09 (2 days prior to the start of group housing) and on GD14 (the end of group housing period) with blood collection day (PD09 and GD14) as a fixed factor and a random slope and subject ID and as a random intercept.

### 2.4 Transcriptomic data analysis

The workflow of transcriptome data analysis is summarized in **Fig. 1B**.

#### 2.4.1 TagSeq data processing

Raw reads were processed to obtain gene count data by following the TagSeq data processing pipeline provided based on Meyer et al. (2011) and Lohman et al. (2016). Briefly, customized perl script utilizing FASTX-Toolkits (Hannon, 2010) CUTADAPT v. 2.8 (Martin, 2011) was used to remove reads with a homo-polymer run of “A”≥8 bases and retain reads with minimum 20 bases and removal PCR duplicates. Processed reads were then mapped to the Mus Musculus reference genome (Ensembl release 99) using Bowtie2 (Langmead and Salzberg, 2012). Quality of raw sequence data was checked with FastQC (Andrews, 2010).

#### 2.4.2 Deconvolution

We used Cell-type Identification By Estimating Relative Subsets Of known RNA Transcripts (CIBERSORT) (Newman et al., 2019) deconvolution algorithm to estimate abundances of immune cell types from RNAseq data. Tissue-specific training datasets for mouse species were obtained from Chen et al. (2018) and uploaded to CIBERSORTx website and the analysis for each tissue was performed with 1000 permutations, relative run mode, disabled quantile normalization and batch correction. With those immune cell types showing high abundance in each tissue, we tested whether cell abundance is different between alpha and subordinate mice by fitting Bayesian regression with social group ID as a random factor.

#### 2.4.3 Differential gene expression (DGE) analysis

Principal component analysis was conducted with filtered gene counts data (filtered out genes with less than 10 counts for each sample) with a voom transform. We used the *limma* R package (Smyth et al., 2021) to identify differentially expressed genes (DEGs) in liver and spleen transcriptome data respectively with i) social status (alpha vs. subordinate) and ii) GD14 plasma corticosterone (CORT) concentrations. Raw p-values are adjusted via empirical false discovery rate (eFDR) (Storey and Tibshirani, 2003). We permuted sample labels for 5000 times and obtained a null distribution of p-values to estimate empirical false discovery rate. We set a threshold for differentially expressed genes as 15% change of the absolute values of log2 fold change at the empirical false discovery rate (eFDR) of 5%. Enriched gene ontology (GO) analysis was conducted to explore functional differences among four different groups of DEGs (upregulated in alphas vs. in subordinates, positively vs. negatively associated with corticosterone). We used the clusterProfiler R package (Yu et al., 2020) to test statistical overlap between annotated biological and functional significance in each DEG set.

#### 2.4.4 Weighted gene co-expression network analysis (WGCNA)

Weighted gene co-expression network analysis was performed using the WGCNA R package (Langfelder and Horvath, 2008). As we collected 11 samples from alpha males and 12 samples from subordinates and the use of WGCNA analysis is recommended to use with a total sample number greater than 20, we aggregated count data across social status groups (total of 23 samples) and constructed one gene co-expression network independently for each tissue transcriptome. Both WGCNA models were constructed in signed hybrid mode with minimum module size set as 30 genes. The power values were set 6 for liver transcriptome and 18 for spleen transcriptome. Gene counts were normalized with the *limma* (Smyth et al., 2021) package and genes that have low expression counts (<50) in more than 90% of samples were filtered out prior to network construction. Based on the scale-free topology criterion (Zhang and Horvath, 2005), we set the power parameter as 5 for liver, 14 for spleen transcriptome to construct a signed hybrid correlation network. Linear mixed effects model was performed with each behavioral or biological measurement as a fixed factor and social group ID as a random factor to determine the association between each module eigengene and other traits of interest. Enriched gene ontology (GO) analysis for genes contained in each module was conducted via the clusterProfiler R package (Yu et al., 2020). For the modules of interest noted on Fig. 3C, we constructed protein-protein interaction (PPI) network based on STRING database which contains known and predicted protein-protein interactions (Szklarczyk et al., 2019) to identify genes playing a critical role in each module. The construction and visualization was performed by the STRINGdb R package (Franceschini, 2021). We entered mouse entrez IDs of genes in each module to GeneWalk software (version 1.5.2) to identify relevant GO terms for genes of interest in each module and further identify regulator genes and moonlighter genes.

#### 2.4.5 Differential co-expression network analysis (CoDiNA)

To investigate genes potentially playing critical roles in a specific condition (alpha or subordinate status), we applied Co-expression Differential Network Analysis (CoDiNA) (Gysi et al., 2020). In the transcriptome of each tissue, gene co-expression networks are constructed with data from alpha males and subordinates separately, calculating a weighted topological overlap (wTO) value for each gene-to-gene connection (link) present in each network (Gysi et al., 2020). Subsequently, we filtered the links with top 1% absolute wTO values and entered them to CoDiNA (liver transcriptome: 0.378 ; spleen: 0.794). CoDiNA identifies alpha links (links that are present in both alpha and subordinate gene networks in the same direction), beta links (present in both conditions but co-expressed in the opposite direction) and gamma links (specific only to one condition and does not exist in the other condition).

## 3. Results

We housed 120 male CD-1 mice in pairs in standard-size cages for 10 days (**Fig. 1A**) and determined their dyadic social status based on aggressive interaction observed between pair housing day (PD) 06-10. Then we assigned mice from each pair to different social groups of ten mice each, counterbalancing the number of pair-dominant and pair-subordinate mice in the newly formed social group. Each mouse social group was housed in a group housing system (**Fig. 1C**) designed to mimic the natural habitat of wild mice (Berry, 1970; Williamson et al., 2016) for 14 days. Trained observers recorded aggressive interactions noted on the ethogram (Supplemental Table S1) every day during group housing periods (median of total observation hours 29.5, interquartile range (IQR) = (28.8 – 30.2)).

### 3.1 All social groups form significantly linear hierarchies

All 12 social groups in the current study formed significantly linear dominance hierarchies (Table S3; Landau’s h value = median 0.89, IQR = (0.82 – 0.93), all p-value < 0.01; triangle transitivity = 0.91 (0.83 – 0.97), all p-value < 0.01). Significantly high directional consistency across all social groups (0.93 (0.92 – 0.95)) indicated that most of the wins were directed from dominant to subordinate males. The despotism of each group varied with a median and IQR of 0.45 (0.42 – 0.60) ranging from a minimum of 0.33 to a maximum of 0.63. **Fig. 1D** & **Fig. 1E** show the win/loss sociomatrix and temporal dynamics of Glicko ratings of each mouse in an exemplar cohort (Cohort I). The sociomatrices and Glicko rating graphs of all cohorts are provided in Supplemental Figure S1 and Supplemental Figure S2. In one cohort (Cohort G) the original alpha male was displaced by the beta male on day 6 of group-housing, but the linearity and stability of the hierarchy were quickly re-established and maintained.

### 3.2 Dominant mice in pairs show higher concentrations of corticosterone than subordinates

We measured concentrations of plasma CORT from all subjects twice at Zeitgeber time 06, on the pair housing Day 9 (PD09) and on the last day of group housing (GD14). In pair housing, dominant mice had significantly higher CORT than subordinate mice (**Fig. 2A**; parameter estimates b_Dom-Sub_: 36 ng/ml, 95% credibility intervals (CI) = [12,59]).

**Figure 2.**
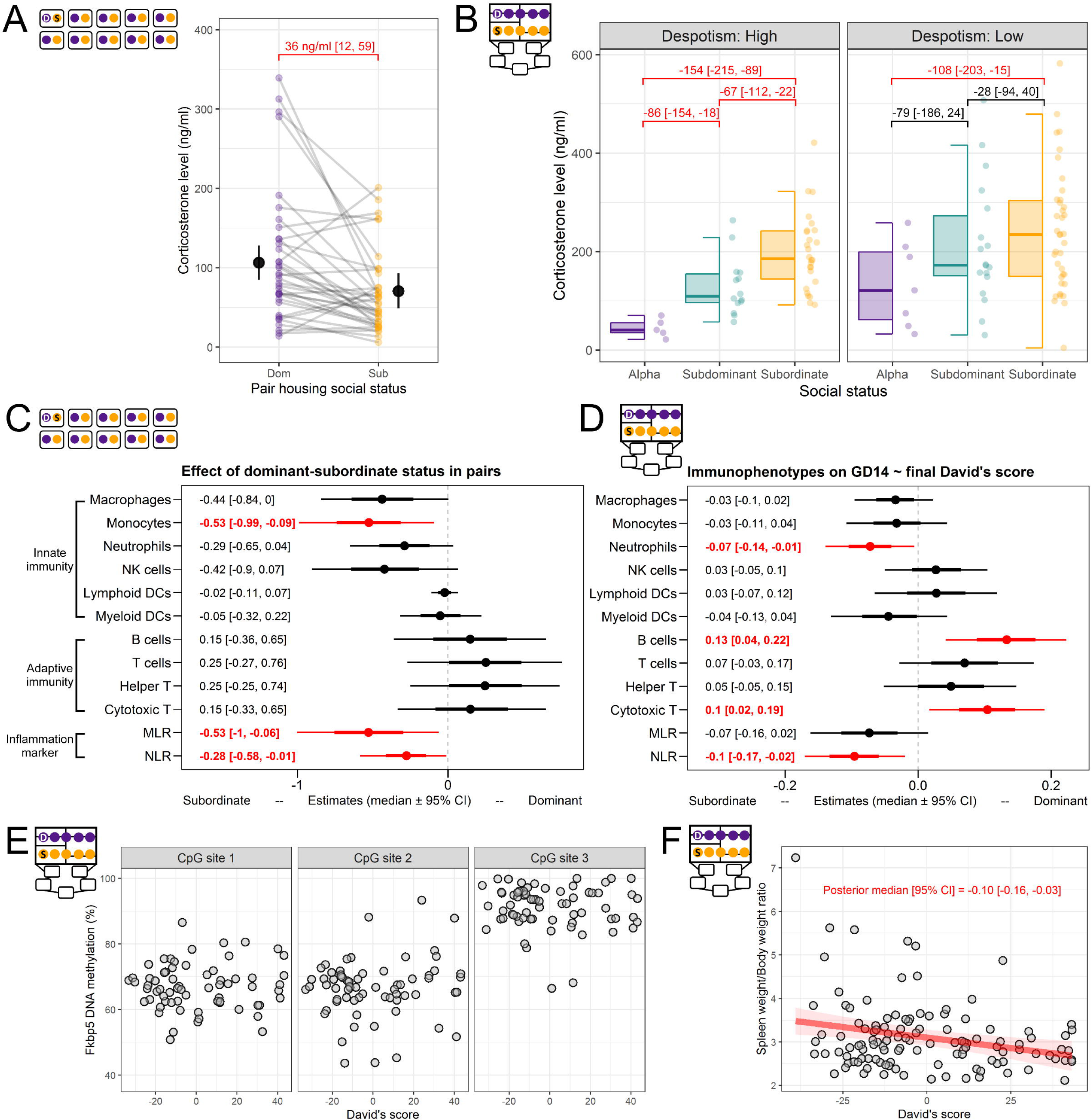
(A, B) Plasma corticosterone levels by social status measured during pair housing (PD09) and after 14 days of social housing (GD14). Immunophenotype differences by dyadic dominant-subordinate status (PD09) (C) and by social dominance score in hierarchies (final David’s score, GD14) (D). The graphs represent parameter estimates and 95% credibility intervals for each Bayesian linear regression analysis for posterior densities, posterior medians (solid dots), 66% credible intervals (thick bars), and 95% credible intervals (thin bars) for comparison. The summary text in A-D notes median estimates and 95% CI for each comparison. (E) *Fkbp5* DNA methylation levels of the three measured CpG sites by David’s score. David’s score was not significantly related to DNA methylation levels in any of the measured CpG sites. (F) Mice with lower David’s score have relatively higher spleen weight (adjusted for body weight).

**Figure 3.**
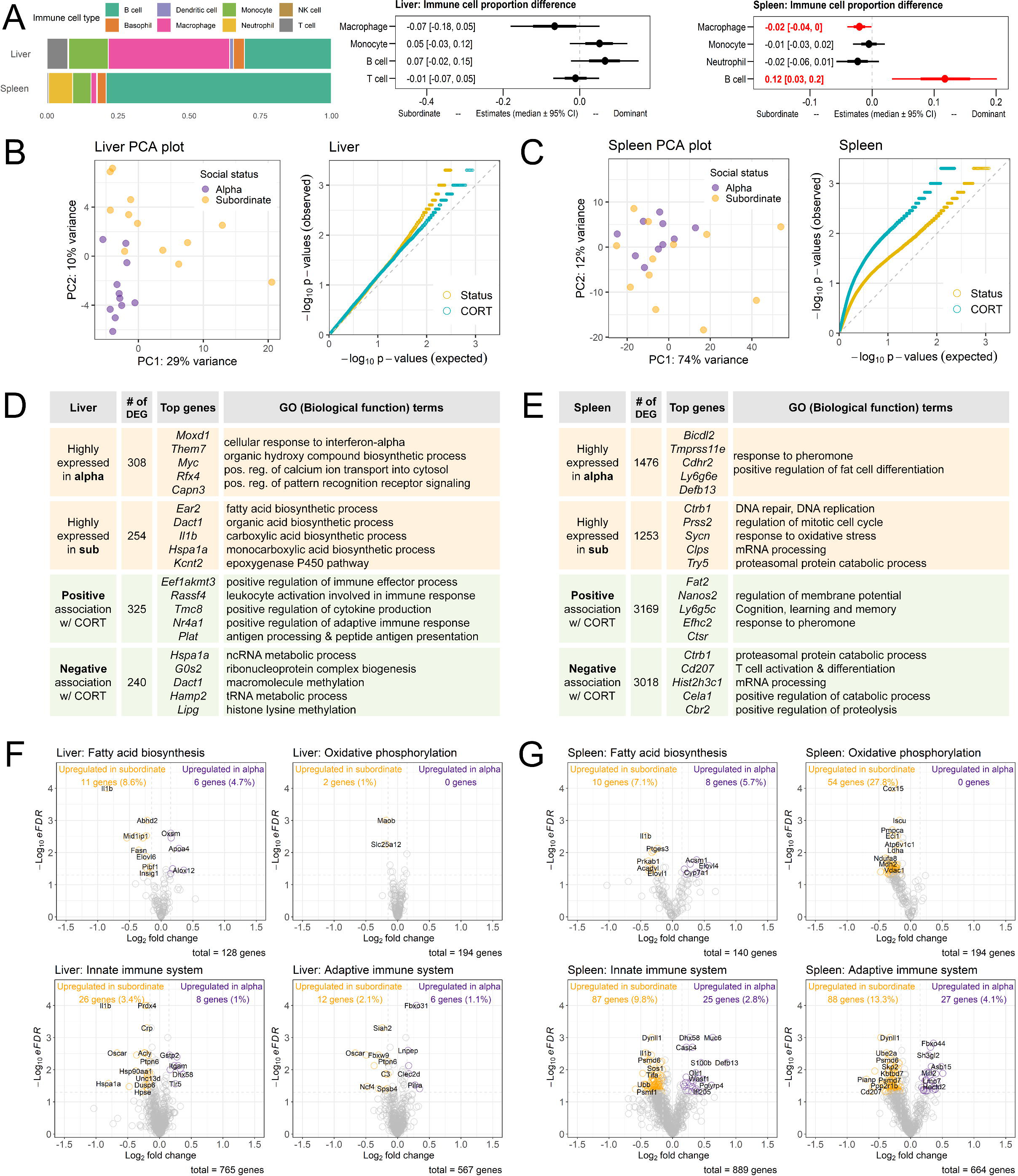
(A) Estimated immune cell abundance and differences in estimated cell proportions of each cell type between alpha and subordinate males in liver and spleen. (B,C) Principal component analysis (PCA) of RNA-seq data. QQ-plots demonstrate the distribution of eFDR values associated with the effects of social status versus corticosterone (GD14) in each tissue. Each comparison is against the expected null (uniform) distribution. (D,E) Summary of DEG analysis by socials status or by corticosterone (GD14) and Gene Ontology Biological Process (GO-BP) analysis for upregulated or downregulated DEGs for each comparison. Top genes represent five genes with the highest absolute value of log2 fold change. (F,G) Hypothesis-driven exploration of DEGs. Volcano plots show DEG effect size and significance (eFDR) for genes in the selected gene sets. Differentially expressed genes (DEGs) met criteria if the absolute values of log2 fold change were greater than 15% at the empirical false discovery rate (eFDR) of 5%.

### 3.3 Subordinate mice in social hierarchies show higher concentrations of corticosterone and the difference is greater in despotic hierarchies

The relationship between CORT and social status was reversed after two weeks of living in social hierarchies. We categorized the 12 social groups into high- and low-despotism hierarchies using the same cut-off value as in previous studies (0.5, within the scale 0 to 1; Williamson et al., 2017). In both high- and low-despotism social groups, alpha males had significantly lower CORT than subordinate males (Fig. 2B; high-despotism: b_alpha-subordinate_ = -154 ng/ml [-215,-89]; low-despotism: b_alpha-subordinate_ = -108 [-203,-15]). In the highly despotic hierarchies, we observed significant differences between alphas and subdominants (b_alpha-subdominant_ = -86 [-154,-18]) and between subdominants and subordinates (b_subdominant-subordinate_ = -67 [-112,-22]). These differences were not significant in the low-despotism hierarchies. These effects of social status on circulating CORT across housing conditions and hierarchy despotism replicate our previous findings (Williamson et al., 2017). Additionally, we tested if the large variation of CORT in the subordinate population in hierarchies (**Fig. 2B**) might be explained by varying levels of received aggression, spleen weights, or pair housing social status. However, we did not find any significant associations.

### 3.4 Subordinate mice in pairs show a higher proportion of monocytes, monocyte:lymphocyte ratio (MLR), and neutrophil:lymphocyte ratio (NLR) in blood

We assessed immunophenotypes of all animals before (PD09) and after (GD14) social hierarchy formation. Using two-panel multicolor flow cytometry, we measured the proportion of immune cells in peripheral blood involved in innate (macrophages, monocytes, neutrophils, natural killer (NK) cells, lymphoid dendritic cells (DCs), myeloid DCs), and adaptive (B cells, cytotoxic and helper T cells) immunity (Unsworth et al., 2016). Additionally, we also analyzed monocyte:lymphocyte ratio (MLR) and neutrophil:lymphocyte ratio (NLR). High values of these ratios indicate inflammation (Mazza et al., 2018; Xiang et al., 2018). Mice that were subordinate in pair housing had significantly higher monocytes (b_Dom-Sub_ = -0.53 [-0.99, -0.09]), and a trend towards more macrophages (b_Dom-Sub_ = -0.44 [-0.84, 0]), neutrophils (b_Dom-Sub_ = -0.29 [-0.65, 0.04]) and NK cells (b_Dom-Sub_ = -0.42 [-0.9, 0.07]) (**Fig. 2C**). Further, subordinate mice had significantly higher MLR and NLR ratios (MLR: b_Dom-Sub_ = -0.53 [-1.00, -0.06]; NLR: b_Dom-Sub_ = -0.28 [-0.58, -0.01]) than dominant mice. These findings are consistent with socially defeated individuals that exhibit an elevated inflammation response (Hodes et al., 2014; Powell et al., 2013).

### 3.5 Macrophage proportion prior to group housing predicts final social status in hierarchies

We tested if cell proportions measured prior to hierarchy formation predicts individual final social rank in hierarchies. Mice with higher levels of macrophages before hierarchy formation were more likely to occupy a lower social position in the hierarchy regardless of their pair-housing dominance status (-0.54 [-1.06, -0.03], Supplemental Figure S3).

### 3.6 Subordinate mice in hierarchies show lower B and cytotoxic T cell % and higher NLR in blood

Social status measured by David’s scores was associated with immunophenotypes measured on GD14 (**Fig. 2D**). More dominant (higher David’s score) males had significantly higher levels of B cells (0.13 [0.04, 0.22]) and cytotoxic T cells (0.10 [0.02, 0.19]) and lower levels of neutrophils (-0.07 [-0.14, -0.01]) than lower-ranked individuals. Subordinate mice also had a trend towards more macrophages (-0.03 [-0.1, 0.02]) and monocytes (-0.03 [-0.11, 0.04]) than higher-ranked mice. Consistent with this, high ranked individuals had significantly higher NLR (-0.10 [-0.17, -0.02]) and a trend towards higher NK cells (0.03 [-0.05, 0.1]) and helper and cytotoxic T cells (0.07 [-0.03, 0.19]) than lower-ranked individuals.

### 3.7 Mice show significant changes in cell proportions after 14 days of group housing

We tested if there are significant changes in the proportion of each cell type after being housed in group housing for 14 days (Supplemental Figure S4). We found there are significant decreases in B lymphocytes (-0.91 [-1.14, -0.68]), cytotoxic T lymphocytes (-0.40 [-0.64, -0.16]), and significant increases in levels of monocytes (0.40 [0.13, 0.66]) and neutrophils (0.79 [0.53, 1.04]), as well as MLR (0.57 [0.29, 0.86]) and NLR (0.81 [0.53, 1.09]) across all animals. While there was not a statistically significant interaction between cell proportion change and social status, B cell % decreased 10.2% in subordinates (29.9 to 19.8%) compared to alphas decreasing 5.3% (31.6 to 26.3%).

### 3.8 DNA methylation levels of the *Fkbp5* gene are not associated with social dominance

Previous studies (Ewald et al., 2014; Kitraki et al., 2015; Lee et al., 2011) suggest that chronic CORT administration is linearly associated with higher *Fkbp5* gene expression and lower *Fkbp5* DNA methylation levels in various tissues including blood cells in mice. We assayed DNA methylation levels across three CpG sites located ∼200bp upstream of Exon 5 of the *Fkbp5* gene as a proxy of chronic circulating CORT concentrations. DNA methylation levels of each CpG site were not associated with either David’s score measure of social dominance (**Fig. 2E**) nor CORT concentrations on GD14 (Supplemental Figure S5).

### 3.9 Spleen weight is associated with social dominance and alpha males with higher despotism show higher spleen weights

Mice with lower David’s score had relatively higher spleen weight (adjusted for body weight, mg/g) (**Fig. 2F**; -0.10 [-0.16, -0.03]). The spleen weight of each alpha male from each social hierarchy was compared against their despotism. Alpha males with higher despotism showed significantly higher spleen weight (Supplemental Figure S6; 2.71 [0.94, 4.55]).

### 3.10 Deconvolution of hepatic and splenic transcriptome reveals differences in the estimated immune cell proportion between alpha and subordinate mice

The spleen and liver perform numerous functions involved in the immune response, including development, storage and blood trafficking of lymphocytes, the production of antibodies, and filtering out and breaking down old or damaged blood cells while harboring their useful components (Carsetti et al., 2004; Jenne and Kubes, 2013; Tarantino et al., 2013). Thus, we isolated total RNA from liver and spleen tissues of the alpha (Rank 1) and the most subordinate (Rank 10) mice in each social group. We used Tag-based RNA sequencing (Tag-Seq; Lohman et al., 2016) and followed standard raw data processing pipelines to obtain raw count data (see Section 2.4 for more details).

To determine abundance of immune cell types in liver and spleen transcriptome data, we used a Cell-type Identification By Estimating Relative Subsets Of known RNA Transcripts (CIBERSORT) (Newman et al., 2019) deconvolution algorithm. Tissue-specific training datasets for mouse species were obtained from Chen et al., (2018). In the hepatic transcriptome, the most abundant cell types were B cells, macrophages, monocytes, and T cells and there were no significant differences between alpha and subordinate mice (**Fig. 3A**). In the splenic transcriptome, the estimated proportion of B cells was significantly higher in alpha male samples compared to those of subordinate males. In contrast, the macrophage proportion was significantly higher in subordinates than in alphas. High levels of innate immune cells in spleen and blood of subordinate males in this study has also been observed in stress-susceptible and socially defeated male mice in social stress paradigms (Ambrée et al., 2018; Avitsur et al., 2002; Bartolomucci et al., 2001; Brachman et al., 2015; Engler et al., 2004; Hodes et al., 2014; McKim et al., 2016; Niraula et al., 2018; Wohleb et al., 2014).

### 3.10 Mice living in social hierarchies show distinct transcriptomic profiles according to social status and plasma corticosterone concentrations

We used the *limma* R package (Smyth et al., 2021) to identify differentially expressed genes (DEGs) in liver and spleen transcriptome data with respect to i) social status (alpha vs. subordinate) and ii) group-housing day 14 (GD14) CORT levels. Differentially expressed genes were considered if the absolute values of log2 fold change were greater than 15% at an empirical false discovery rate (eFDR) of 5% (see Section 2.4.3 for details).

Principal component analysis (PCA) revealed distinct separation between alpha and subordinate males in the liver transcriptome (**Fig. 3B**), whereas PCA of the spleen transcriptome was less separated by social status (**Fig. 3C**). There was no distinct pattern in the separation of PCAs of liver and spleen transcriptome according to GD14 CORT concentrations (Supplemental Figure S7). The QQ-plots (**Fig. 3B**) comparing the eFDR distribution associated with social status and CORT indicates that in the liver transcriptome the social status effect was stronger than the effect of CORT. The opposite pattern was observed in the splenic transcriptome. In the liver, among the 13100 genes we tested in *limma* model (filter counts >20 in 90% of samples), 562 (4.3%) genes were differentially expressed by social status and 565 (4.3%) genes were identified as DEGs associated with CORT (**Fig. 3D**). Notably, we observed a total of 24 *Mup* genes that were detected in the liver transcriptome, and 88% of those genes (21 genes) were significantly upregulated in alpha males compared to the subordinates (Supplemental Figure. S8). In the splenic transcriptome, among 17515 genes tested in DEG analysis, social status was associated with differential expression of 2729 (15.6%) genes and 6187 (35.3%) genes were differentially expressed by CORT (**Fig. 3E**). Interestingly, a significant number of genes encoding olfactory receptors (*Olfr*, 677 genes) and vomeronasal receptors (*Vnmr*, 204 genes) were observed. Among those, 20% (136 genes) of the identified *Olfr* genes and 30% (62 genes) of the identified *Vnmr* genes were identified as DEG, and all of those genes except for one gene (*Vnm2r66*, downregulated in alpha) were upregulated in alpha males.

**Fig. 3D** and **Fig. 3E** summarize the results of DEG analysis and corresponding Gene Ontology Biological Process (GO-BP) analysis for both tissue samples. Interestingly, multiple biosynthetic processes were enriched in those genes upregulated in liver tissue of subordinates. These biosynthesis processes have in common that they require energy, such as in the form of Adenosine triphosphate (ATP) or nicotinamide adenine dinucleotide phosphate (NADPH) (Barger and Plas, 2010). DEGs positively associated with CORT were enriched in the immune effector process, leukocyte activation, adaptive immune response, and cytokine production. In the spleen, subordinate animals exhibited an up-regulation of genes involved in DNA replication and repair and mitotic cell cycle regulation. DEGs negatively associated with CORT in the spleen were enriched in T cell activation and differentiation. We further explored patterns of gene regulation across both tissues. A total of 206 genes were significantly differentially expressed by social status in both tissues and 23% of those genes (47 genes) were expressed in the opposite direction. A total of 391 genes were significantly associated with CORT in liver and spleen. 30% of those genes (119 genes) were regulated in the opposite direction by CORT. For example, the *Nr4a1* gene, which has one of the largest positive associations with CORT in the liver and is involved in glucocorticoid receptor binding and regulation of autophagy and apoptosis (Muller, 2017), is negatively associated with CORT in the spleen transcriptome.

### 3.11 Hypothesis-driven gene set exploration

While GO analysis for up- or down-regulated genes is useful for exploring the functionality of the DEG genes, it is also important to explore the degree and pattern of transcription of genes of interest chosen by pre-existing hypotheses. We explored gene expression patterns of specific gene sets associated with immune systems (innate vs. adaptive, and cytokine signaling) and energy metabolism (fatty acid biosynthesis, oxidative phosphorylation, glycolysis) (**Fig. 3F**, **Fig. 3G**; Supplemental Figure S9). Counter to our expectations, there was no difference in the number of genes differentially upregulated by social status in the four selected gene sets in the liver transcriptome (**Fig. 3F**). In the splenic transcriptome, we observed large differences in numbers of genes upregulated by status in each of the curated gene sets. A striking difference was observed in the oxidative phosphorylation gene set, in which all DEGs (28%) were upregulated in subordinate males (**Fig. 3G**). Similarly, the genes involved in glycolysis were more upregulated in subordinate males (12%) compared to in alpha males (3%). In both innate and adaptive immune system gene sets, the number of the genes upregulated in subordinates were significantly higher than the number of genes upregulated in alpha males (alpha males: innate 3%, adaptive 4%; subordinates: innate 10%, adaptive 13%). Additionally, based on the previously identified GO term (**Fig. 3E**), we confirmed that 16% of genes in the DNA repair gene set were upregulated in subordinates and 2% were upregulated in alphas.

### 3.12 Weighted gene co-expression network analysis reveals discrete gene networks associated with social status and other biological phenotypes

We further explored the Tag-Seq data with Weighted Gene Correlation Network Analysis (WGCNA). WGCNA identifies clusters of genes that behave similarly across samples (Langfelder and Horvath, 2012, 2008). We constructed a WGCNA network for each tissue type including both alpha and subordinate samples in each network. After identifying clusters of co-expressed genes (modules) within each network, we can calculate module eigengene (ME), the first principal component of gene expression profiles of all genes contained in the module (Langfelder and Horvath, 2008). We tested each ME for the association with behavioral and biological traits measured in the study using mixed-effect linear regression (LMM) analysis (social group ID as a random factor). Heatmaps (**Fig. 4A** and **Fig. 4B**) show beta coefficients from the LMM for statistically significant associations between MEs and traits.

**Figure 4.**
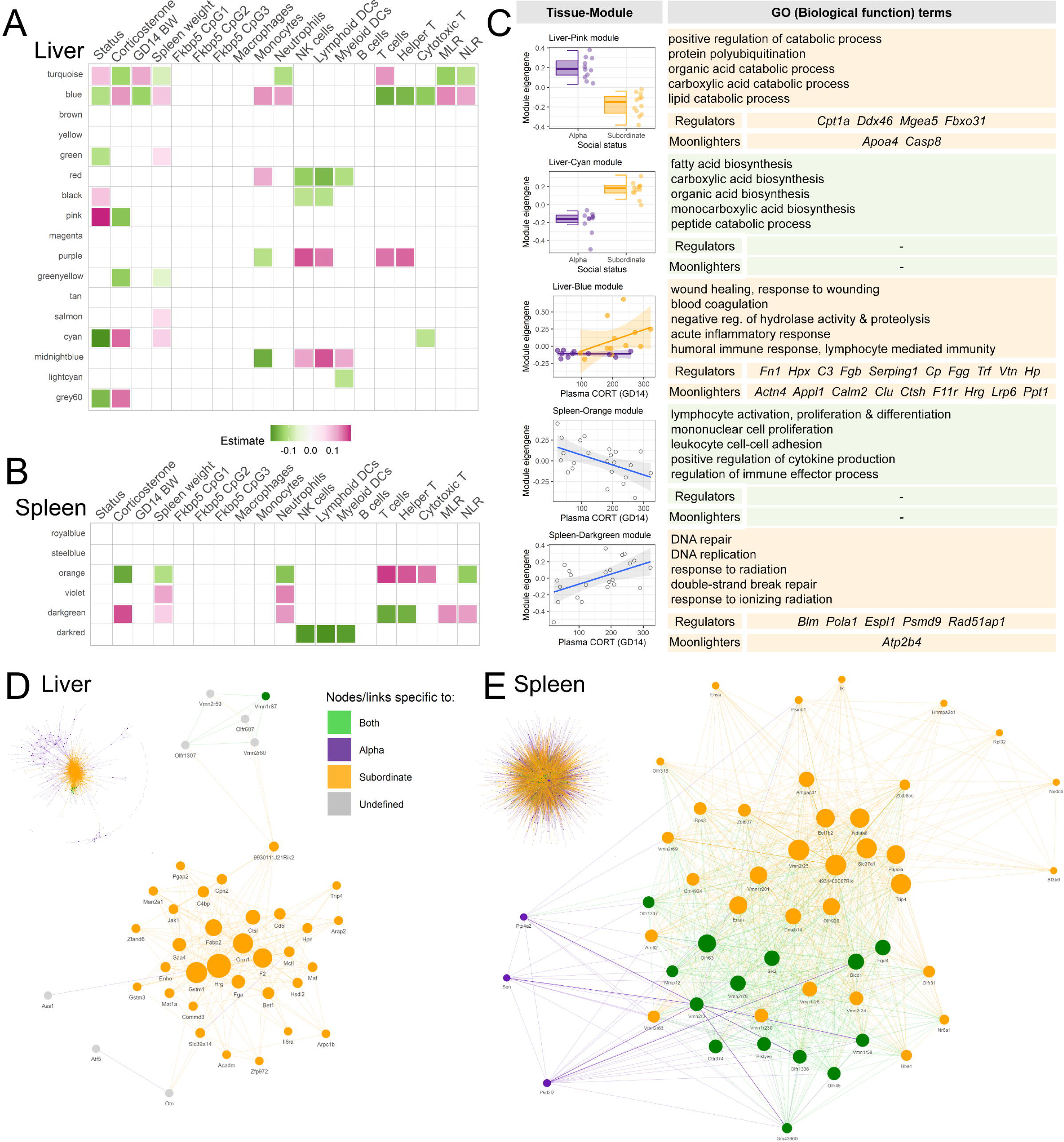
Results of Weighted Gene Correlation Network Analysis (WGCNA) and Co-expression Differential Network Analysis (CoDiNA). (A,B) Heatmaps representing linear regression estimates of eigengene expression of WGCNA modules against other measured traits. Only significant associations (p-value <0.05) are represented. (C) Module eigengene expression by social status or corticosterone (GD14) and summary of GO-BP analysis and GeneWalk analysis for five selected modules. (D,E) CoDiNA networks. Small network graphs represent links with the strongest co-expression (wTO). To explore meaningful gene networks differentially connected according to social status, we further filtered and visualized 50 gene nodes with the highest degree centrality.

In the liver transcriptome, WGCNA identified a total of 17 modules (**Fig. 4A**). Among those modules, the eigengenes of the pink and cyan modules showed striking differences between alpha and subordinate males with no overlap between data distributions being observed. The pink module contained genes upregulated in alpha males, including 19 *Mup* genes. Indeed, the five most differentially expressed genes in the pink module were all *Mup* genes (*Mup16, Mup17, Mup18, Mup9, Mup11*; see Supplemental Table S4 for the list of top five DEGs in the selected modules by status or by CORT). Notably, the protein-protein interaction (PPI) network of genes in the pink module (Supplemental Figure S10) suggests that *Apoa4* and *Cyp8b1* genes are closely related to *Mup* gene expression. GO enrichment analysis (**Fig. 4C**, see Supplemental Table S5 for the list of top 10 GO terms for all modules) revealed that pink module genes are enriched for catabolic processes of lipid, carboxylic acid, and organic acid that generally produce energy stored in the form of ATP or NADPH. In contrast, the cyan module genes, which are up-regulated in subordinate males, were enriched with genes involved in biosynthetic processes. The blue module was associated with social status and multiple biological measures including CORT. Notably, ME of the blue module is positively associated with CORT only in subordinate males. The blue module genes were enriched for biological function GO terms of wound healing, response to wounding and blood coagulation.

In the spleen transcriptome, a total of six modules were identified by WGCNA (**Fig. 4B**). Interestingly, none of the MEs were significantly associated with social status. Rather, two modules, orange and darkgreen, represented genes that are associated with CORT (**Fig. 4C**). The ME of the orange module was negatively associated with CORT and the genes in this module were enriched for various immune functions, such as activation, proliferation and differentiation of lymphocytes and monocytes and cytokine production. The darkgreen ME was positively associated with CORT and represents genes involved in DNA replication and repair.

We further investigated genes in each WGCNA module with GeneWalk (**Fig. 4C**). GeneWalk assembles a network of GO terms from a given set of genes and prunes the network to contain GO terms only specific to an experimental condition. With this approach, GeneWalk is able to curate GO terms only relevant to the given experimental condition among all GO terms registered to each gene in a knowledge-based database (Ietswaart et al., 2021). GeneWalk also identifies regulators and moonlighters. Genewalk identifies regulators as i) genes with high connectivity to other genes and ii) the majority of registered GO terms of those genes are identified relevant to the given GO term network; these transcriptional regulators are likely inducers of transcriptional changes in other genes of their module. We highlight regulators with highest intramodular connectivity (kIN) in **Fig. 4C**. Moonlighters are those genes that have high numbers of registered GO terms but only a low fraction of their GO terms were identified as relevant via GeneWalk, suggesting that their role in the given condition in the study might be specific to specific functions.

### 3.13 Differential co-expression network analysis (CoDiNA) reveals *Hrg* as a central node in the subordinate-specific gene network in liver transcriptome

We implemented Co-expression Differential Network Analysis (CoDiNA) (Gysi et al., 2020) in addition to WGCNA in effort to find converging evidence on potential key driver genes specific to social status. Briefly, a gene co-expression network is constructed for each condition of interest (i.e., a separate network for alpha vs. subordinate males), then CoDiNA identifies co-expression patterns that are: i) present in both alpha and subordinate gene networks in the same direction, ii) are present in both networks but expressed in the opposite direction, iii) or are specific only to one network. We retained the top 1% highly connected links for alpha and subordinate gene networks for differential co-expression analysis (see Section 2.4.5 for details). Both in liver and spleen CoDiNA analysis, most of the links and gene nodes were present only in either the alpha or subordinate network (**Fig. 4D** and **Fig. 4E**; 99% in liver , 94% in spleen). The majority of these links were only present in the subordinate transcriptome (92% of the links in liver, 84% of links in spleen), indicating that the subordinate specific gene networks are more strongly interconnected than alpha networks. Consistent with this, the majority of nodes (genes) in the subordinate gene networks were specific to those networks (52% in liver, 57% in spleen). Conversely, fewer nodes (genes) in the alpha networks were specific to those networks (12% in liver, 6% in spleen). **Fig. 4D** represents a fraction of the liver CoDiNA network (filtered to contain top 50 genes with the highest degree centrality). Notably, among genes identified as moonlighters in the Liver-blue module from WGCNA and GeneWalk, *Hrg* (a gene for histidine rich glycoprotein) is also the central node of the differential co-expression network identified by CoDiNA. As *Hrg* is a specifically subordinate network gene node, this suggests that *Hrg* and other genes closely co-expressed with *Hrg* are driving transcriptional adaptation to social status in the liver.

## 4. Discussion

Variations in social experience are associated with changes in immune system functioning but the directionality and magnitude of these relationships appear to be highly context-specific. Our findings extend previous evidence to demonstrate that specific factors related to status in male mice living in social hierarchies, namely social stress exposure and energy requirements, shape immune and metabolic strategies.

Examination of immune cell populations in peripheral blood and RNA-seq analysis of liver and spleen revealed status-related variation in innate versus adaptive immunity. Innate immunity, which constitutes the first line of defense against pathogens and is metabolically more expensive to maintain than antigen-specific adaptive immunity (Habig and Archie, 2015; Lee, 2006), appear relatively more abundant in blood and splenic tissue of subordinate male mice compared to that of dominant male mice in both pair-housed and group-housed contexts. The relatively low levels of energetically expensive innate immune cells in dominant males compared to subordinate males is consistent with the life history trade-off hypothesis that dominant animals direct most of their energy away from immune processes and towards other physiological processes related to maintaining their high status (Habig and Archie, 2015; Lee, 2006). Accordingly, we found that peripheral blood and splenic tissue of group-housed, but not pair-housed, dominant mice contain relatively higher proportions of cells involved in the energetically cheap adaptive immune response that is designed to defend against specific acute stressors that are potentially more common in the lives of high-ranking animals (Habig and Archie, 2015; Lee, 2006; Sapolsky, 2004). Specifically, we found that they had greater proportions of B lymphocytes and cytotoxic T cells in peripheral blood and a greater estimated proportion of B cells in their splenic transcriptome. Differential B cell concentration in the spleen convincingly reflects differential adaptive immune function in dominant and subordinate animals, as the spleen is responsible for the maturation, storage, and trafficking of B cells (Carsetti et al., 2004; Tarantino et al., 2013) to the extent that splenectomy in mice results in complete absence of the mobile B cell subtype in circulating blood (Carsetti et al., 2004). While our current knowledge of how social experiences shape immune function in dominant animals is limited, our results are broadly consistent with prior studies that have shown that high-ranking female rhesus macaques have greater levels of cytotoxic T cells (Tung et al., 2012) and high-ranking female baboons have greater levels of B cells (Lea et al., 2018) in peripheral blood than low-ranking individuals.

Spleen weight is also negatively associated with social status in hierarchies of Brandt’s voles (Li et al., 2007), rats (Blanchard et al., 1985; Spencer et al., 1996), and mice (Turney and Harmsen, 1984), including mice subjected to social defeat stress (Foertsch et al., 2017; McKim et al., 2018). While physical injury is the most common explanation for immune cell trafficking and increased weight in the spleen (Foertsch and Reber, 2020; Turney and Harmsen, 1984), the subordinate mice in this study did not exhibit overt signs of wounding. They do, however, receive a disproportionate amount of aggression from higher-ranking males, which can result in sympathetic hyperactivity and consequently increased spleen weight (Wehle et al., 1978). Similarly, mice who are repeatedly defeated in social stress paradigms do not always acquire physical injuries and unwounded animals exhibit immune cell responses that follow, albeit with reduced magnitude, the same trend as wounded animals (Engler et al., 2004). Indeed, exposure to stressors ranging from physical to psychosocial in nature results in noradrenergic stimulation in the spleen and local upregulation of myelopoiesis and enthropoiesis, accumulation of neutrophils and monocytes, and consequently increases in volume and mass of the spleen (Dahlstroem and Zetterstroem, 1965; Foertsch et al., 2017; McKim et al., 2018). Interestingly, we also found that amongst group-housed alpha males, spleen weight was positively associated with despotism. Alpha males rarely receive attacks once the hierarchies are formed, and therefore increased spleen weight is likely not due to bite wounds but rather the stress associated with having to consistently assert aggression to maintain their status (Williamson et al., 2016), thus supporting the idea that spleen weight may reflect levels of psychosocial as well as physical stress.

Interestingly, we did not find significantly higher proportions of adaptive immune cells in the peripheral blood of pair-housed dominant animals. A likely explanation for this finding is that the energetic demands of being dominant in a group and in a semi-natural environment are much greater than being dominant in a dyadic context. In fact, all group-housed animals exhibited elevated levels of innate immune cells and decreased levels of adaptive immune cells compared to those in pair-housing regardless of status, suggesting that group-housing is a chronically and generally stressful environment. It is also important to note that across social ranks, the group-housing context was more predictive of the animals’ immunophenotype. In other words, an animal’s social experience prior to group-housing did not appear to influence their immune cell composition, such that status-related differences in innate versus adaptive immune cell proportions emerged even in cases where animals changed their social status after pair-housing. This is somewhat contrary to findings in female rhesus monkeys whose plasma immune profiles are affected by both current and previous rank, although current rank is more predictive (Sanz et al., 2020; Tung et al., 2012). Also, in male baboon hierarchies, immune gene profiles appear to predetermine social rank more than they are shaped by social rank (Lea et al., 2018). This suggests that species and sex-specific social factors mediating the formation and maintenance of dominance hierarchies likely differentially shape how immunophenotypes are influenced by current versus previous social status.

There is accumulating evidence that social status can profoundly impact transcriptomic profiles across central and peripheral tissues including blood in primates (Lea et al., 2018; Snyder-Mackler et al., 2016; Tung et al., 2012) and brains of mice (Fujita and Yamashita, 2021; Saul et al., 2019, 2017), zebrafish (Oliveira et al., 2016), sticklebacks (Greenwood and Peichel, 2015; Saul et al., 2019), and honey bees (Saul et al., 2019). Here we extend these findings by demonstrating that male mice living in a social hierarchy showed distinct hepatic and splenic transcriptomic profiles according to both social status and CORT. Gene expression PCA plots for the liver showed almost complete separation between alpha and subordinate males which was not as robust with the splenic transcriptome. Although social status did influence differential splenic gene expression, transcriptomic profiles of spleen were more affected by CORT than status. This distinction is important as the relationship between circulating CORT and social status is not straightforward (Williamson et al., 2017). Group-housed subordinate mice did show higher CORT compared to dominant males, but this effect was only large in individuals living in despotic hierarchies. Further, we did not find any significant effect of social status on *Fkpb5* DNA methylation. This probably suggests that animals of different ranks are not experiencing sufficient variations in CORT exposure over the two weeks of housing to induce differences in DNA methylation of this gene. Thus, while it is likely that subordinate animals do experience some increase in CORT exposure compared to dominant males in social hierarchies, there is also considerable individual variation, indicating that not all status-phenotype associations are driven by changes in HPA signaling and that circulating CORT can also induce phenotypic changes independently of rank.

It is well established that elevated levels of stress including social stress can lead to increased HPA and sympathetic nervous system activity that induce apoptosis of lymphocytes causing reduced lymphocyte cell numbers and suppressed lymphocyte function (Gurfein et al., 2014; Jiang et al., 2017; Lucin et al., 2009; Stevenson et al., 1989). Consistent with this, using WGNCA we identified a co-expressed module (orange) of splenic genes involved in lymphocyte activation, proliferation and differentiation including *Cd10, Tnfrsf13b, and Tmem131l,* whose activity were negatively associated with CORT. Using a hypothesis-driven approach, we explored the effect of social status on expression patterns of innate and adaptive immune function gene sets. Consistent with our hypothesis, we observed that subordinate males had more upregulated DEGs in the innate immunity gene set in the splenic transcriptome. Contrary to our expectations however, subordinates also had a higher upregulation of DEGs in the adaptive immunity gene set compared to alpha males. This suggests that the reduced numbers of lymphocytes observed in subordinates may be as a result of inhibition of lymphocyte production rather than a proliferation of these cells in dominants. Supporting this, we identified up-regulation of the *Tgfb* gene in subordinates (Supplemental Figure S9), which encodes for the cytokine TGF-β which inhibits B-cell proliferation and modifies T-cell differentiation. We hypothesize that accumulated exposure to subordination stress in social hierarchies leads to differential expression of *Tgfb* as well as CORT induced reduced expression of genes promoting lymphocyte proliferation indicated by the orange module that ultimately lead to lower splenic and blood levels of B cells and cytotoxic T cells (Jiang et al., 2017; Wang et al., 2007).

Long-term exposure to social stress and CORT can potentially have severe deleterious effects for the health of tissue. Using WGCNA we identified a module (dark-green) of co-expressed genes involved in DNA replication and repair whose expression was highly positively associated with CORT exposure. DEG and GO analysis (**Fig. 3E**) also identified that the most up-regulated genes in the spleens of subordinates are involved in mitotic cell cycle regulation and subsequent DNA replication and repair. For instance, 81 DEGs related to DNA repair including *Spire2, Ube2a, Btg2* and *Hmces* showed significantly higher expression in subordinates compared to only 12 DEGs showing higher expression in dominants (Supplemental Figure S9). We also observed a striking difference in splenic expression in the oxidative phosphorylation gene set, in which all 54 DEGs were upregulated in subordinate males. These include subunits of NADH dehydrogenase such as *Ndufa8* and *Nudfv1* that are part of the electron transport chain in mitochondria and *Iscu* and *Pmpca* that regulate enzymatic functioning in mitochondria. It is possible that spleen cells are utilizing oxidative phosphorylation to generate the significant cellular energy required to support DNA repair and ameliorate tissue damage (Brace et al., 2016).

Hepatic transcription profiles are even more strongly influenced by social status than those in the spleen. Notably, our data demonstrate several genes associated with inflammation and liver disease are differentially expressed by status. Subordinate males show upregulation of *Il1B* the gene encoding for the proinflammatory cytokine interleukin 1 beta (IL-1B) that is a major mediator of proinflammatory responses, whereas alpha males barely showed any expression. IL-1B is barely detectable in healthy livers, but is expressed by activated monocytes and macrophages following infection, liver injury or other insults such as exposure to drugs or toxins (Tilg et al., 2016; Tsutsui et al., 2015). Persistent activation of IL-1B leads to various liver diseases such as cirrhosis, fibrosis, and non-alcoholic steatohepatitis (NASH) (Miura et al., 2010; Tilg et al., 2016). One of the immune processes involving IL-1B signaling is the activation of inflammasomes. However, we did not find significant transcription level differences in any of the genes involved in IL-1B processing and NLRP3 inflammasome activation, such as *Il18, Nlrp3, Asc, Aim2,* and *Casp1* (Guo et al., 2015; Syed et al., 2020). This suggests potential status-dependent health outcomes in liver are mediated by other downstream IL-1B signaling pathways. The oncogene *Myc* is upregulated in alpha males and encodes the transcription factor MYC. MYC is well known for its role in coordinating the transcriptional regulation of genes that promote a multitude of cellular functions including cell-cycle regulation, cell survival, cell proliferation, metabolism, and protein and ribosomal synthesis (Miller et al., 2017). MYC can influence these processes in both adaptive and innate immune cells (Gnanaprakasam and Wang, 2017). Notably, however, MYC has been found to regulate the expression of immune checkpoint genes including CD46 and programmed death-ligand 1 (Casey et al., 2018). This suggests that high expression of *Myc* in alpha males may inhibit highly proliferative cells from eliciting their immune response.

WGCNA and GO analysis also identified a module (blue) in the liver that included co-expressed genes related to blood coagulation, wound healing processes, inflammatory and immune responses that are up-regulated in subordinate males, particularly in those with elevated CORT. Included in these genes, are 6 out of 8 coagulation factor genes (*F2, F5, F7, F9, F10, F12*) expressed in the liver. *Fga* (fibrinogen alpha-chain) promotes the synthesis of the alpha portion of fibrinogen and is upregulated in response to tissue injury to help clot formation and induce inflammation (Gabay and Kushner, 1999; Ryu et al., 2015). *Hrg* is highly connected with the other genes in the module and its protein facilitates blood coagulation, activates lymphocytes and supports healthy liver functioning (Guo et al., 2020; Johnson et al., 2014; Poon et al., 2011). Shifts in the expression of these genes in subordinates with high CORT is likely to occur in response to accumulating injuries in subordinates. While we do not observe detrimental levels of physical injury from the socially housed mice, accumulation of minor injuries may be sufficient to invoke such liver transcriptome changes. Alternatively, alpha males may be forgoing investment in these processes as a trade-off to direct energy investment into other more critical processes.

Alpha and subordinate males living in social hierarchies have divergent energetic demands. Correspondingly, social status was associated with strikingly divergent expression patterns of metabolism related genes in the liver. WGCNA revealed a module (pink) of co-expressed genes that were enriched for organic acid, carboxylic and lipid catabolic processes that were up-regulated in dominants. Conversely, a second module (cyan) of co-expressed genes were up-regulated in subordinates and were enriched for carboxylic acid, monocarboxylic acid, organic acid and fatty acid biosynthesis as well as peptide catabolism. This suggests a dramatic shift in metabolism, with dominant animals keeping up with high energy demands by breaking down lipids and oxidizing fatty acids to release energy while inhibiting their synthesis.

Specifically, we found that alpha males had significantly higher expression of the *PNPLA2* gene which encodes the adipose TG lipase (ATGL) enzyme. ATGL facilitates the hydrolysis of triacylglycerols to diacylglycerols and is the first and rate limiting step of lipolysis (Zechner et al., 2012). Alpha males also show drastically lower expression of *G0s2* which encodes the lipolytic inhibitor G0/G1 switch protein 2 (G0S2). G0S2 is a negative inhibitor of ATGL, and thus a reduction in expression leads to increased expression of ATGL. Indeed, G0S2 is considered to be a master regulator of storage of fatty acids in adipose tissue versus mobilization in the liver (Zhang et al., 2017). Further, we observed an up-regulation in alpha males of several genes in the pink module promoting fatty acid oxidation. The solute carrier protein *SLC27A2* is responsible for transportation of free fatty acids from blood into the cytosol of liver cells (Anderson and Stahl, 2013). Carnitine palmitoyltransferase 1A (CPT1A) is a rate limiting step enzyme that connects long-chain fatty acids to carnitine thus enabling its transfer from cytosol across mitochondrial membranes into mitochondria where fatty acid oxidation occurs (Kersten, 2014). *SLC22A5* encodes the protein OCTN2 which transports carnitine into the cell to facilitate this transfer of free fatty acids into mitochondria (Longo et al., 2016) and *Ces2c* encodes the carboxylesterase Acylcarnitine hydrolase that catalyzes O-acylcarnitine to produce available carnitine (Longo et al., 2016). Clearly, alpha male mice have altered expression of genes at all levels of the fatty acid oxidation pathway to increase energy production. Interestingly, these same pathways are altered in dominant and subordinate rainbow trout but in opposite directions, suggesting that they are highly inducible by social status but in a context specific manner (Kostyniuk et al., 2018).

Mitochondria that are actively engaged in fatty acid oxidation are unable to synthesize fatty acids through negative feedback mechanisms. Congruent with this, expression of key genes that promote fatty acid synthesis such as Fatty acid synthase *Fasn* and acetoacetyl-CoA synethetase, *Aacs* were downregulated in alpha males. Notably, activation of these two genes in immune cells is required for cholesterol synthesis, the activation of macrophages and the production of IL1β (Carroll et al., 2018). Other differentially expressed genes in the co-expression modules were also identified to associate with both metabolism and immune response. ALOX12 is downregulated in alpha males and encodes for arachidonic acid 12-lipoxygenase which catalyzes the oxidation of polyunsaturated fatty acids including arachidonic acid that promotes inflammation responses and wound healing (Tallima and El Ridi, 2017; Zheng et al., 2020). Further, using CoDiNA analyses we also identified that subordinates have a specific co-expressed network of metabolic and inflammatory response genes that was not present in alpha males. Central to this network were the genes *Hrg* and *Orm1* (Alpha-1-acid glycoprotein 1) both of which are known to integrate inflammatory and metabolic signals to shape immune responses (Bartneck et al., 2016; Y. S. Lee et al., 2010), as well as two genes of the glutathione S-transferase (GST) family *Gstm1* and *Gstm3* which encode for the primary phase II metabolic enzymes that promote clearance of reactive oxygen species from cells. Taken together, these findings suggest that subordinate animals likely shift their metabolic activity including moving from fatty acid oxidation to synthesis in part to support their pro-inflammatory responses.

Alpha males living in mouse social hierarchies are faced with a multitude of energetic demands. In addition to increasing their levels of aggressive and patrolling behavior (Williamson et al., 2016), they also greatly increase their production of MUPs in the liver which are then deposited via urine scent-marks and used to communicate their dominance status to other animals (Guo et al., 2015; Lee et al., 2017; Nelson et al., 2015). We have previously demonstrated that dominant animals significantly increase their feeding and drinking compared to subordinate animals to support the production of these signals (Lee et al., 2018). Despite their higher caloric intake, it is notable that their transcriptomic profile is more similar to fasting animals that switch to using fatty acids as an energy source for the liver (Sokolović et al., 2008). This demonstrates just how energetically demanding it is for alpha males to maintain their social status. In the current study, we found that almost all Mup genes, including *Mup20* the most potent indicator of social status, were among the most differentially expressed genes being extremely more highly expressed in the livers of alpha males. The majority of these genes were co-expressed in the pink module, although *Mup3*, *Mup20* and *Mup21* were co-expressed in the turquoise module that was also up-regulated in alpha males. Analysis of the protein-protein interaction (PPI) network for the pink module identified the *Apoa4* gene which encodes for the Apolipoprotein A-IV protein and the *Cyp8b1* gene, a cytochrome P450 protein involved in the production of bile, as both being up-regulated in alpha males and highly interconnected with Mup proteins suggesting a possible role for these genes in the regulation of Mup gene expression. Further, increased protein assembly in alpha males’ livers may also be supported by the expression of *Moxd1* which is upregulated in alphas with a large log fold change. The monooxygenase, DBH-like 1 (Moxd1) protein is a monooxygenase which potentially functions to facilitate inter-conversion of amino acids during periods of amino acid deficiency (Sahar et al., 2011). Additionally, dominant animals may have elevated neuronal demands for energy (Hollis et al., 2018; Larrieu et al., 2017). It has recently been proposed that free fatty acids are an effective source of energy for mitochondria in neurons and rats that are fed medium-chain triglycerides show increased social dominance (Hollis et al., 2018). Potentially, alpha males also increase fatty acid oxidation to support brain mitochondrial functioning to promote their aggressive behavior and maintain their status.

## 5. Conclusion

Our results support the idea that high and low ranking individuals favor adaptive and innate immunity, respectively (Habig and Archie, 2015; Lee, 2006; Snyder-Mackler et al., 2016). In accordance with the life-history trade-off hypothesis, dominant mice exhibit greater investment in energetically cheap adaptive immune components and other processes related to maintaining dominant status, such as the production of MUPs used for territory defense. We also observed upregulation of genes involved in fatty acid oxidation and inhibition of synthesis in the liver of dominant mice, suggesting that these mice are maintaining extremely high energy demands. We also observed higher levels of innate immune cells in peripheral blood of subordinate males compared to dominant individuals and upregulation of splenic genes related to supporting this energetically expensive innate immunity. Consistent with the stress response framework, we did observe higher CORT in subordinates living in hierarchies, which was associated with changes in immune gene expression, such as an upregulation of genes related to wound healing in the liver and DNA repair in the spleen. In conclusion, this study delineates the relationship between social experience and immune function as being dependent on energetic and stress-related pressures. High or low social status does not necessarily predestine an animal to have generally negative or positive health outcomes, but rather an animal’s metabolism and immune function adapt to the pressures of one’s social environment such that certain immune defenses may come at the expense of others.

## Data and Code Accessibility

All raw data and code used in the analyses included in this study are available at the following public repository: https://github.com/jalapic/mouse_socialhierarchy_immune

