## Supplemental Figures for "Distinct inflammatory and transcriptomic profiles in dominant versus subordinate males in mouse social hierarchies"

**Supplemental Figure S1.** Raw sociomatrices of wins and losses for each cohort. Individuals are ordered by I & SI rank. Each value represents the total number of wins by the individual in each row against the individual in each column. The degree of redness represents the frequency of wins.

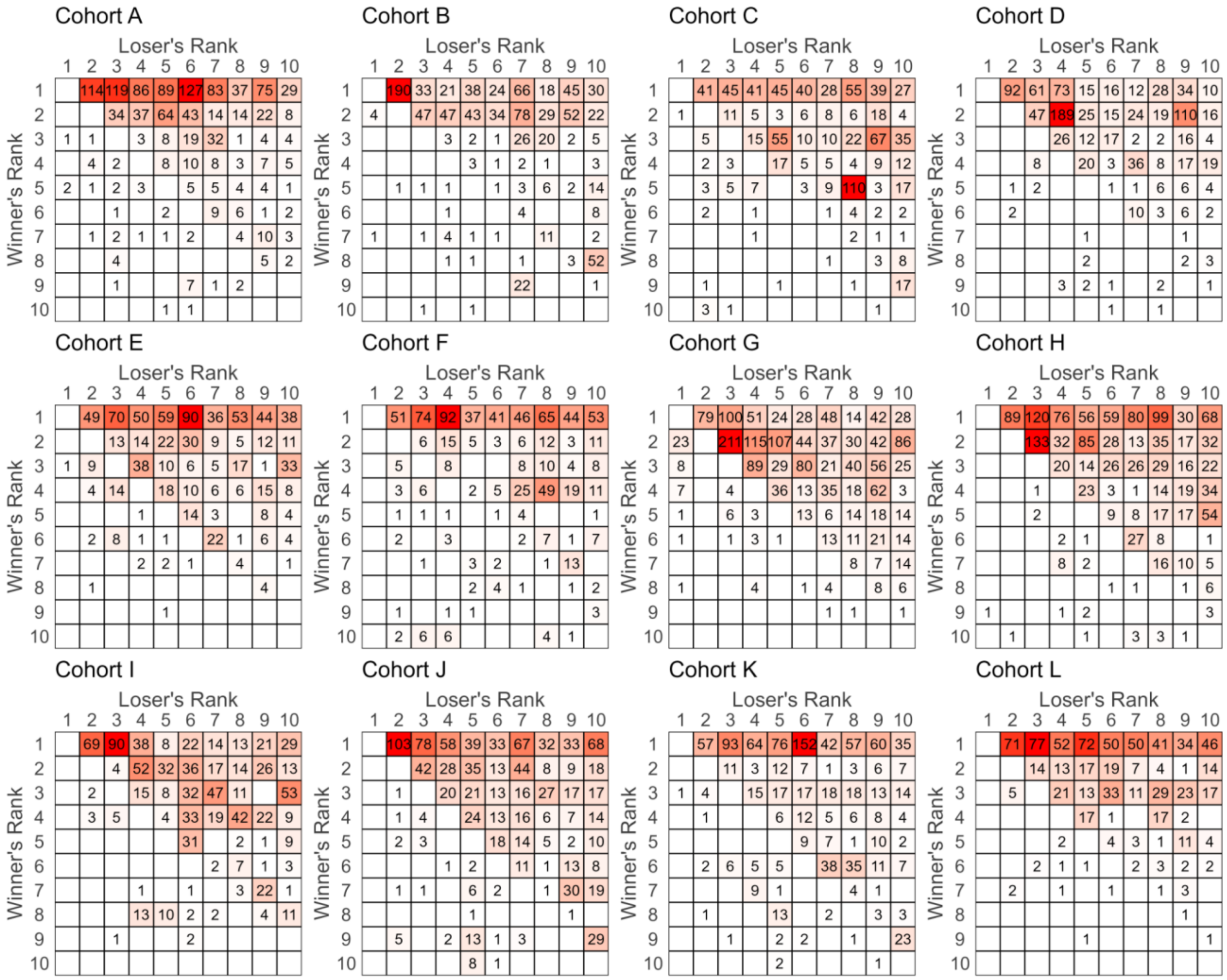

**Supplemental Figure S2.** The Glicko Rating of dominance (y-axis) against each successive behavioral event (x-axis). Each line represents the dominance rating of one individual in each cohort. The lines are colored from black (eventual most dominant individual) to red (eventual least dominant individual).

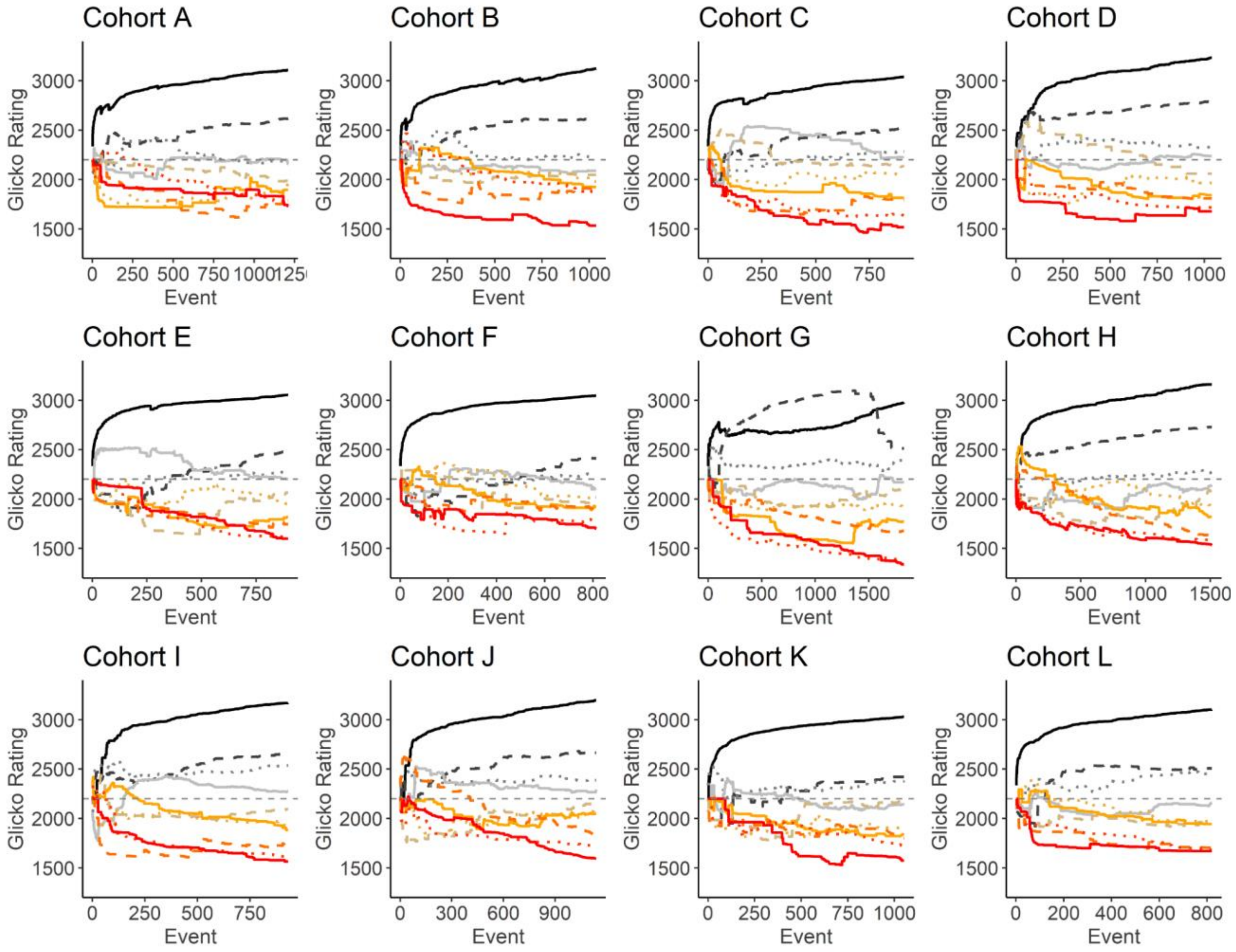

**Supplemental Figure S3.** Estimate of each immune cell level measured prior to social group housing predicting final social dominance score (David's score). Posterior densities, posterior medians (solid dots), 66% credible intervals (thick bars) and 95% credible intervals (thin bars) for comparison among social status groups in each brain region. Text on the left notes median estimates and 95% CI for each comparison.

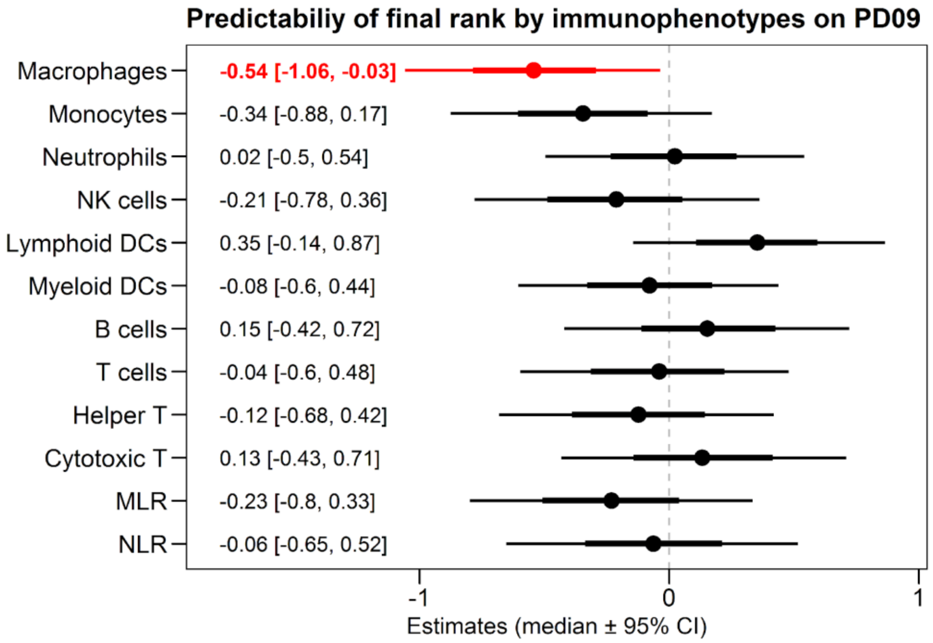

**Supplemental Figure S4.** Changes in the proportion of each cell type from PD09 to GD14. Grey lines represent cell proportion on each time point from the same individual mice. Texts located on the top of the plot indicate parameter estimates and 95% CI. Color labeled rhombi and lines represent mean value of each cluster group and shaded areas represent standard errors of means.

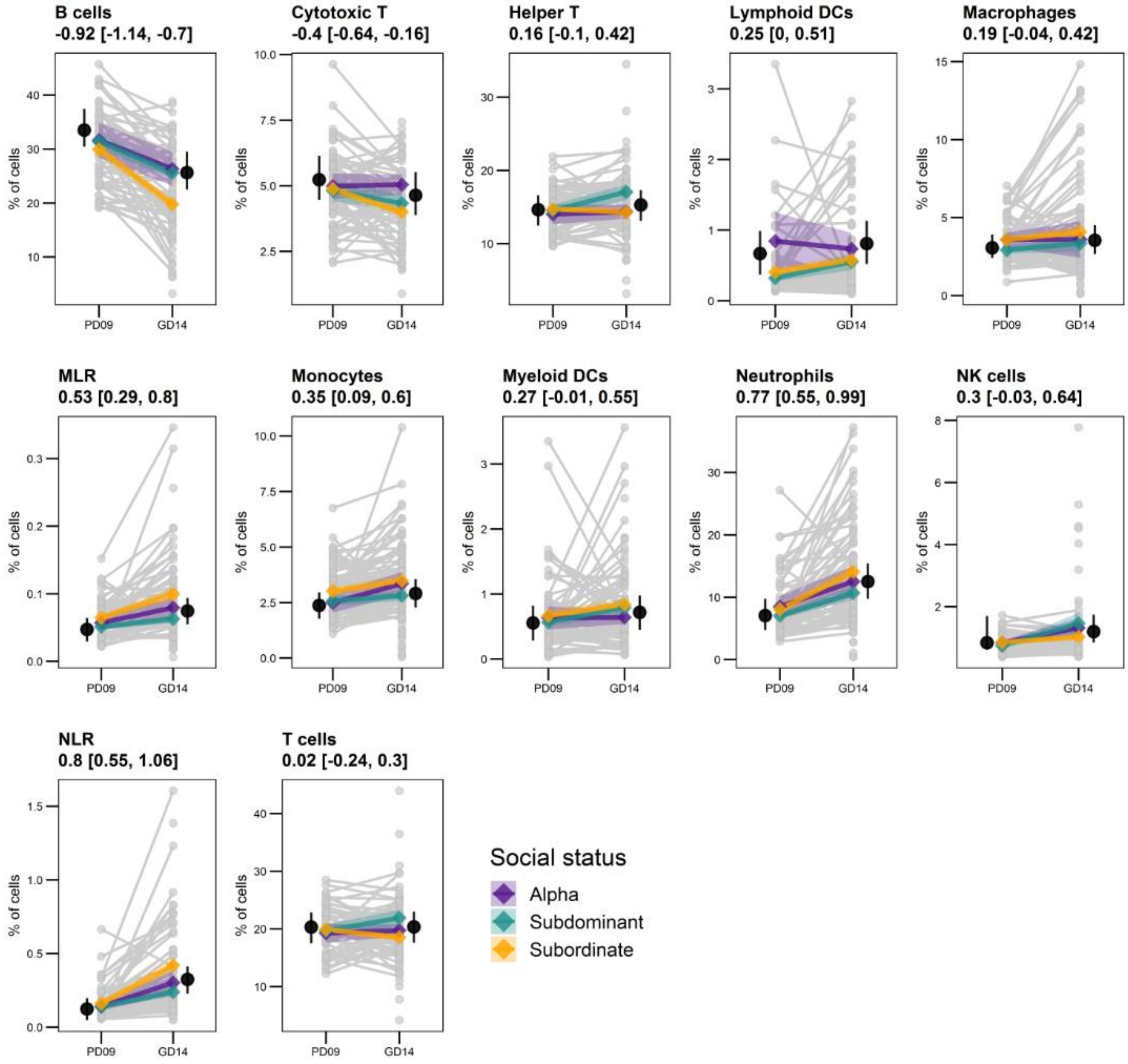

**Supplemental Figure S5.** *Fkbp5* DNA methylation level of the three measured CpG sites by plasma corticosterone levels measured on GD14. There was no significant association between corticosterone levels and the DNA methylation levels in any of the measured CpG sites.

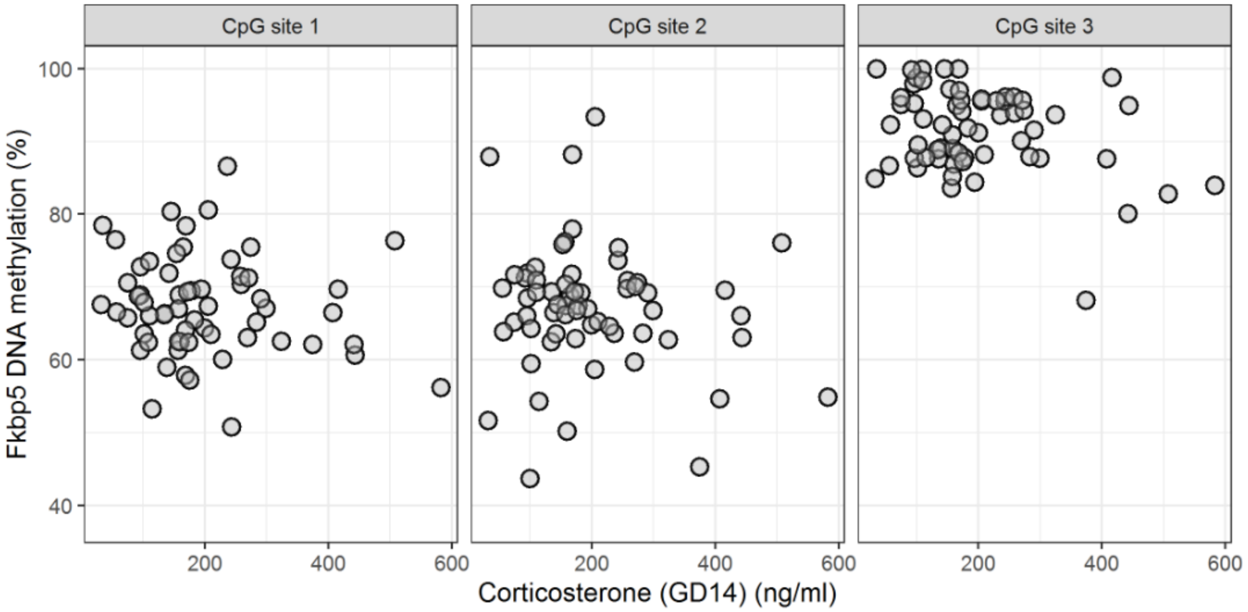


**Supplemental Figure S6.** The spleen weight of each alpha male from each social hierarchy was compared against their despotism (proportion of aggressive encounters won by the alpha males to total aggressive interaction observed in each social hierarchy). Alpha males with higher despotism showed significantly higher spleen weight.

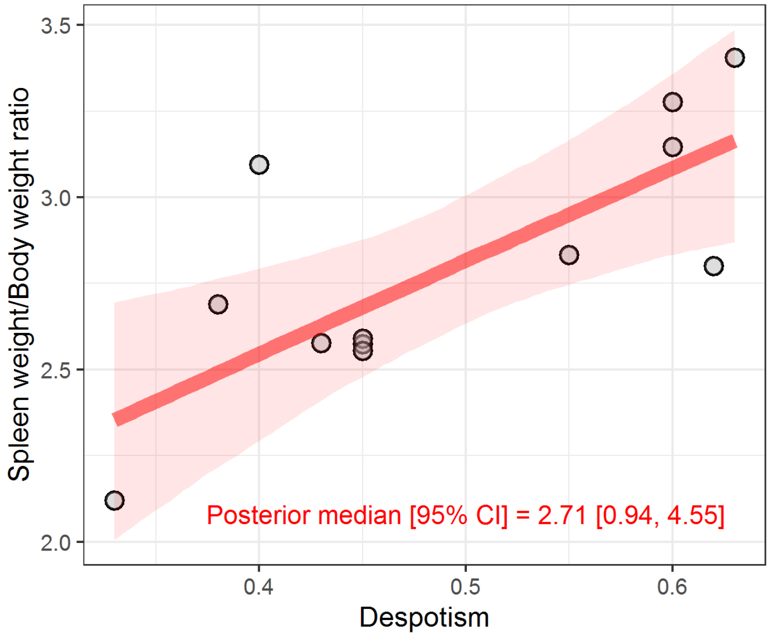

**Supplemental Figure S7.** PCA plot of A) liver and B) spleen transcriptomes labeled by social status and plasma corticosterone levels (GD14).

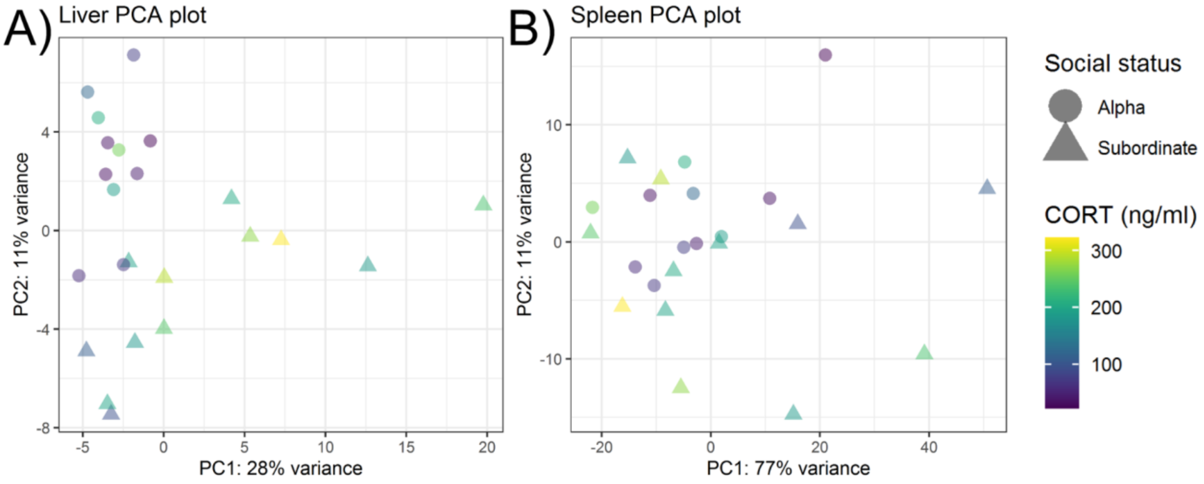


**Supplemental Figure S8.** Normalized gene expression by social status for all *Mup* genes identified in the liver transcriptome.

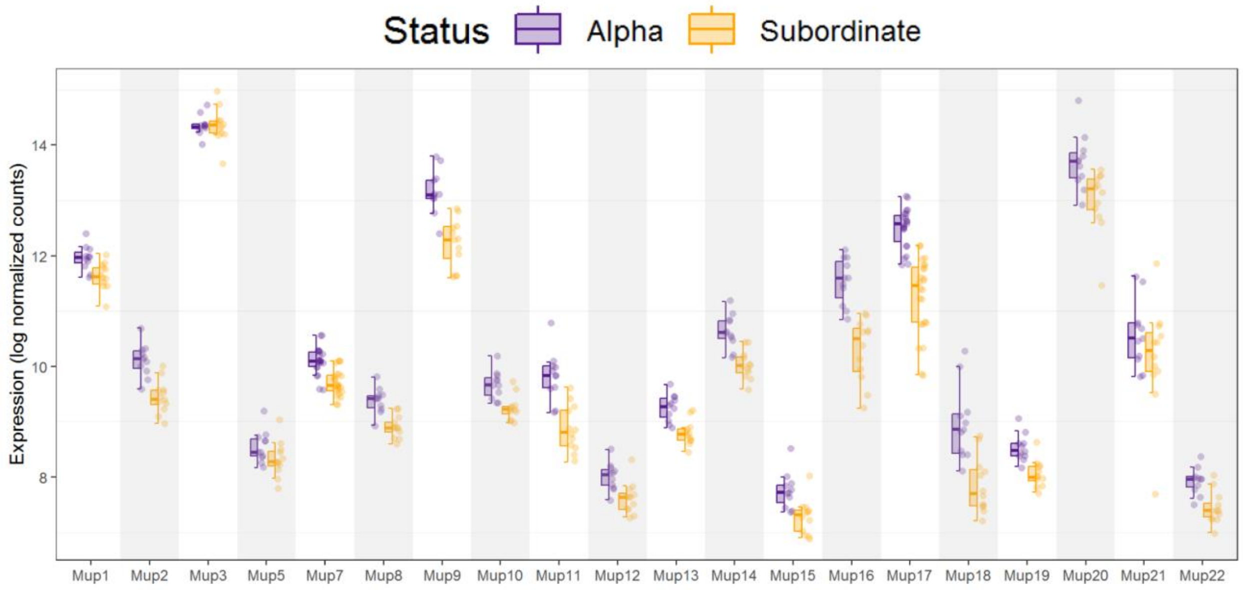

**Supplemental Figure S9.** Volcano plots (DEG by social status or plasma corticosterone levels measured on GD14) for selected gene sets

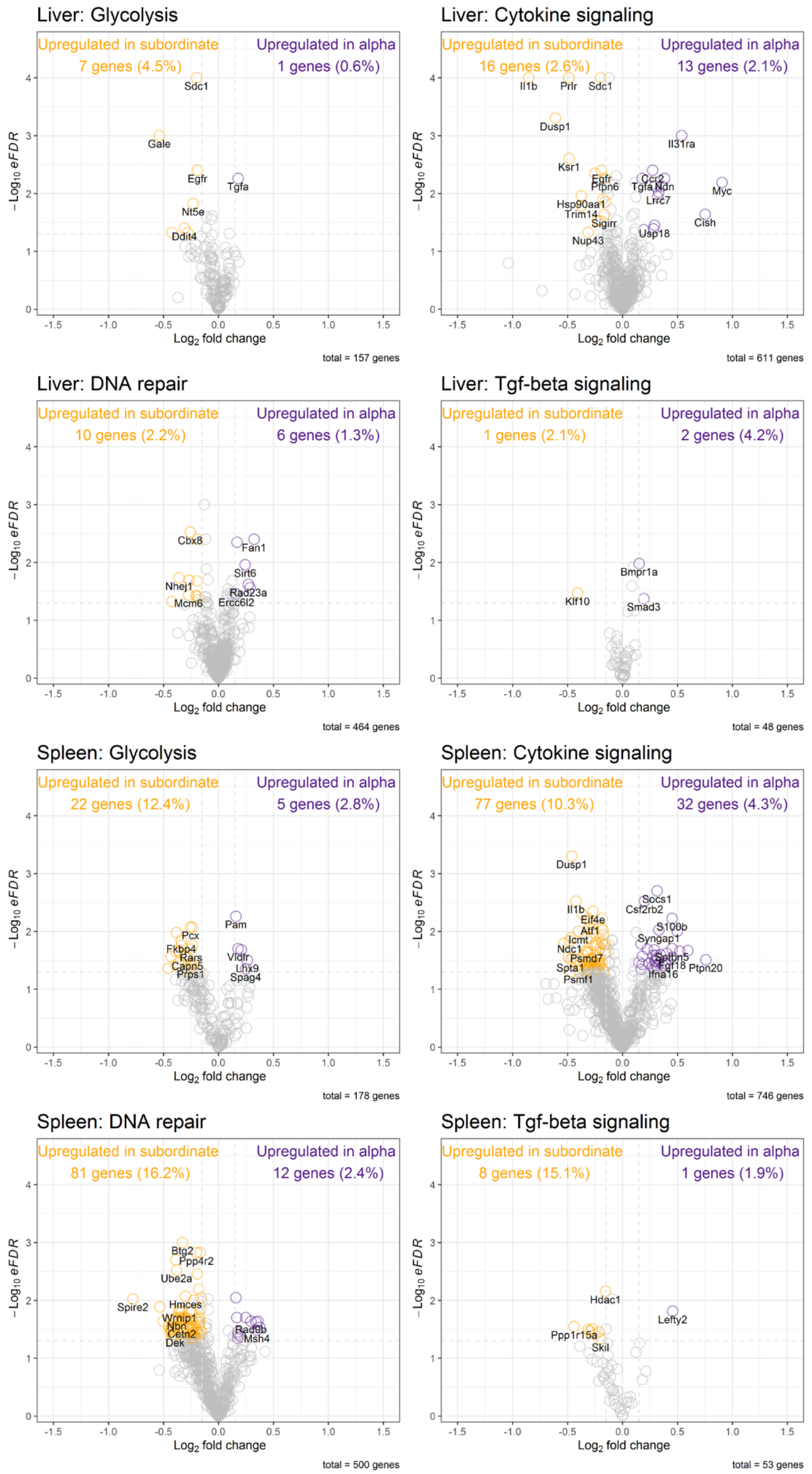

**Supplemental Figure S10**. Protein-protein interaction (PPI) network of pink module genes constructed by STRING

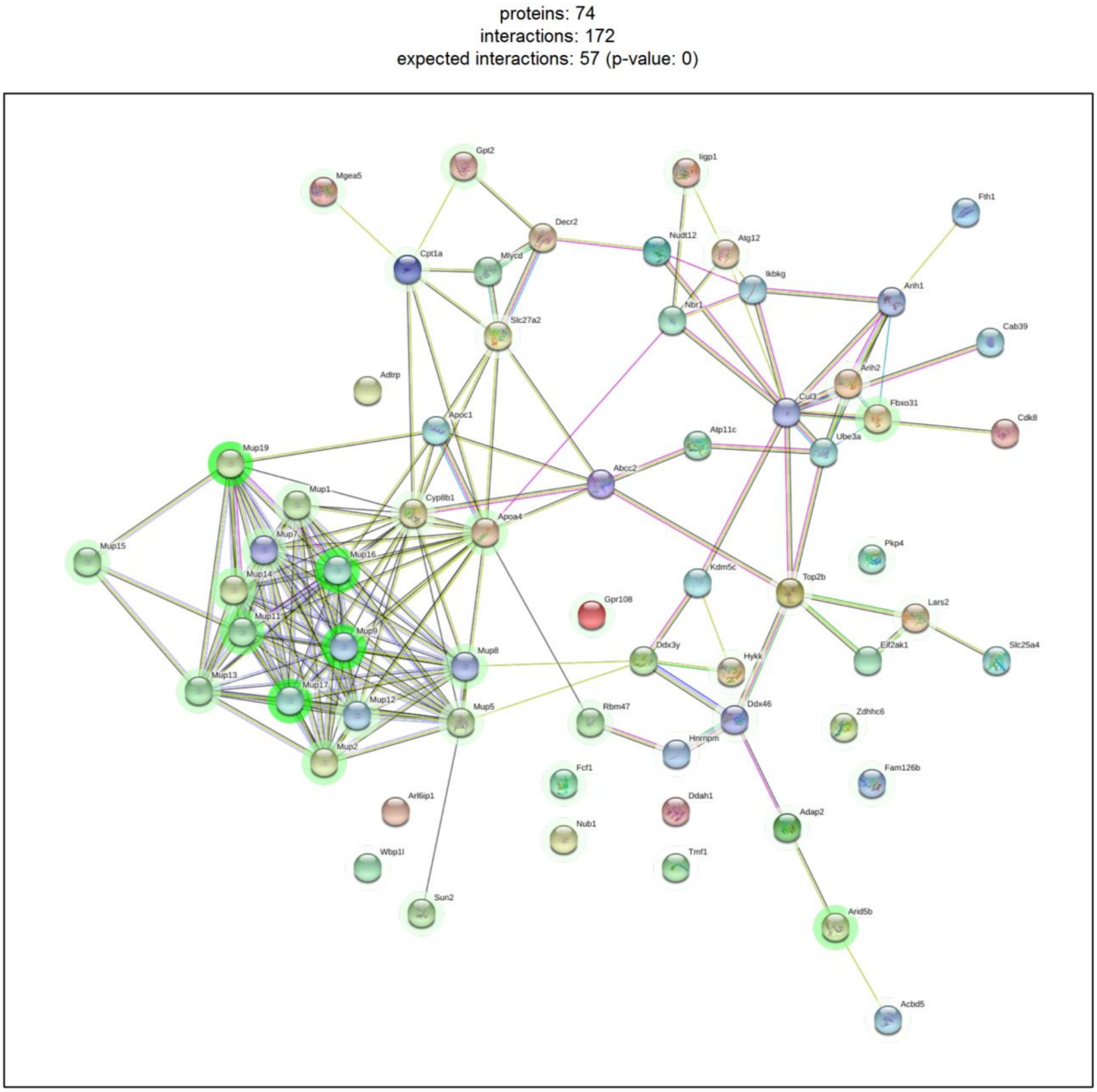
