## Supplemental Tables for "Distinct inflammatory and transcriptomic profiles in dominant versus subordinate males in mouse social hierarchies"

**Supplemental Table S1.** Revised mouse social behavior ethogram (*) Behaviors where one of the two focal animals does not necessarily show any particular behavioral response to the other's behavior.

| State | Behavior | Description |
|---|---|---|
| **Aggressive Behaviors** | Lunge | Mouse lunges an attack at partner with or without making contact to partner |
| | Bite/Fighting | Mouse bites partner on any part of body<br>Individual lunges at and/or bites the other individual |
| | Chase | Individual follows the target individual rapidly and aggressively while the other individual attempts to flee |
| | Mount | Individual mounts another individual from behind with the recipient attempting to flee or otherwise being pinned to the floor |
| **Subordinate Behaviors** | Preemptive avoidance/flee | Without attack or chase from the opponent, mouse rapidly moves away from the opponent as it recognize the presence of the opponent (*) |
| | Flee | Mouse moves rapidly to create maximal distance between themselves and partner |
| | Subordinate posture | Mouse rears head and exposes nape and flank to partner while submitting |
| | Freeze/ Immobility | Mouse stays complete immobile in presence of partner typically also showing piloerection |
| **Other behavior** | Sniff | Mouse olfactory investigates a partner (head/body/anogenital, etc) with or without direct contact (*) |
| | Allogroom | Mouse grooms partner on their fur on any part of their body with mouth or forepaws (*) |

**Supplemental Table S2**. Antibodies used in two-panel multicolor immunophenotyping. Adapted and modified from Unsworth et al. (2016).

| Panel | Antibody | Clone | Fluorochrome | Final dilution | Purpose | Vendor |
|---|---|---|---|---|---|---|
| Panel 1 | CD45.2 | 104 | APC-Cy7 | 1:50 | Leukocytes | Biolegend |
| | CD11c | HL3 | APC | 1:50 | Dendritic cells | BD Biosciences |
| | Ly6G | 1A8 | PE-Cy7 | 1:200 | Neutrophils | BD Biosciences |
| | CD11b | M1/70 | PE | 1:200 | Myeloid cells | BD Biosciences |
| | MHC II | M5/114.15.2 | BV605 | 1:200 | Dendritic cells | BD Biosciences |
| | Ly6C | AL-21 | BV421 | 1:100 | Monocytes, macrophages | BD Biosciences |
| Panel 2 | CD4 | GK1.5 | APC-Cy7 | 1:200 | T-helper cells | BD Biosciences |
| | TCR-beta | H57-597 | PE-Cy7 | 1:200 | T-cells | BD Biosciences |
| | CD45.2 | 104 | PE | 1:50 | Leukocytes | BD Biosciences |
| | CD49b | DX5 | FITC | 1:50 | NK cells | BD Biosciences |
| | CD8 | 53-6.7 | BV605 | 1:200 | Cytotoxic T-cells | BD Biosciences |
| | CD19 | 1D3 | APC | 1:200 | B-cells | BD Biosciences |
| | NKp46 | 29A1.4 | BV421 | 1:100 | NK cells | BD Biosciences |
| Both | - | - | PI | 1:100 | Viability | Biolegend |

**Supplemental Table S3.** Social hierarchy measures for each cohort (A – L).

| | h’ values (p-value) | DC (p-value) | Steepness (p-value) | ttri (p-value) | Despotism | Gini. win | Gini lose |
|---|---|---|---|---|---|---|---|
| A | 0.79 (0.000) | 0.94 (0.000) | 0.71 (0.000) | 0.77 (0.000) | 0.63 | 0.72 | 0.27 |
| B | 0.84 (0.000) | 0.96 (0.000) | 0.69 (0.000) | 0.91 (0.000) | 0.45 | 0.67 | 0.29 |
| C | 0.97 (0.000) | 0.91 (0.000) | 0.80 (0.000) | 1.00 (0.000) | 0.40 | 0.62 | 0.32 |
| D | 0.94 (0.000) | 0.95 (0.000) | 0.74 (0.000) | 1.00 (0.000) | 0.43 | 0.68 | 0.38 |
| E | 0.90 (0.000) | 0.90 (0.000) | 0.67 (0.000) | 0.91 (0.000) | 0.55 | 0.67 | 0.21 |
| F | 0.73 (0.001) | 0.87 (0.000) | 0.57 (0.000) | 0.65 (0.002) | 0.62 | 0.69 | 0.27 |
| G | 1.00 (0.000) | 0.92 (0.000) | 0.90 (0.000) | 1.00 (0.000) | 0.38 | 0.61 | 0.25 |
| H | 0.93 (0.000) | 0.97 (0.000) | 0.82 (0.000) | 0.93 (0.000) | 0.45 | 0.63 | 0.26 |
| I | 0.78 (0.000) | 0.93 (0.000) | 0.73 (0.000) | 0.73 (0.000) | 0.33 | 0.56 | 0.23 |
| J | 0.92 (0.000) | 0.92 (0.000) | 0.76 (0.000) | 0.89 (0.000) | 0.45 | 0.57 | 0.23 |
| K | 0.83 (0.000) | 0.92 (0.000) | 0.73 (0.000) | 0.85 (0.000) | 0.60 | 0.66 | 0.23 |
| L | 0.87 (0.000) | 0.96 (0.000) | 0.69 (0.000) | 0.96 (0.000) | 0.60 | 0.74 | 0.18 |
| Median (IQR) | 0.88 (0.82, 0.93) | 0.92 (0.92, 0.95) | 0.73 (0.69, 0.77) | 0.91 (0.83, 0.97) | 0.45 (0.42, 0.60) | 0.66 (0.62, 0.68) | 0.26 (0.23, 0.28) |

**Supplemental Table S4**. Top 5 DEG by status or by CORT for the selected modules.

| DEG by | module | symbol | logFC | eFDR |
|---|---|---|---|---|
| **Status** | pink | *Mup16* | 0.689516 | 0 |
| | | *Mup17* | 0.673213 | 0 |
| | | *Mup18* | 0.55058 | 0 |
| | | *Mup9* | 0.524857 | 0 |
| | | *Mup11* | 0.485489 | 0 |
| | cyan | *Scd1* | -0.53982 | 0.0035 |
| | | *Ppp1r3c* | -0.43227 | 0.0085 |
| | | *Efna1* | -0.33758 | 0.038 |
| | | *Srebf1* | -0.26528 | 0.0095 |
| | | *Acly* | -0.23132 | 0.003 |
| | blue | *Plcl2* | -0.6875 | 0.023 |
| | | *Ncf4* | -0.47583 | 0.0335 |
| | | *Apcs* | -0.47562 | 0.0115 |
| | | *Hsd17b13* | -0.42453 | 0.0375 |
| | | *Hsp90aa1* | -0.3732 | 0.011 |
| **CORT** | blue | *Hsd17b13* | -0.42634 | 0.034 |
| | | *Hsp90aa1* | -0.41093 | 0.005 |
| | | *Parvg* | 0.343311 | 0.044 |
| | | *Cd44* | 0.324292 | 0.009 |
| | | *S100a6* | 0.301184 | 0.0275 |
| | orange | *AC125374.2* | 0.57593 | 0.005 |
| | | *Mme (Cd10)* | 0.552494 | 0.033 |
| | | *Olfr804* | 0.547099 | 0.007 |
| | | *AC157570.1* | 0.532764 | 0.009 |
| | | *Wdr18* | 0.514863 | 0.023 |
| | darkgreen | *Etnppl* | 0.579668 | 0.0075 |
| | | *Uroc1* | 0.559287 | 0.007 |
| | | *Mgam* | 0.516033 | 0.012 |
| | | *Slc2a9* | 0.44055 | 0.0035 |
| | | *Kcnn2* | 0.433239 | 0.0245 |

**Supplemental Table S5**. Top 10 GO terms for all WGCNA modules not listed in Figure 4.

| module | ID | Description | GeneRatio | p.adjust |
|---|---|---|---|---|
| **green** | GO:0044282 | small molecule catabolic process | 0.104167 | 7.09E-12 |
| | GO:0055086 | nucleobase-containing small molecule metabolic process | 0.1 | 5.36E-07 |
| | GO:0009117 | nucleotide metabolic process | 0.0875 | 2.54E-06 |
| | GO:0006753 | nucleoside phosphate metabolic process | 0.0875 | 2.57E-06 |
| | GO:0072521 | purine-containing compound metabolic process | 0.083333 | 2.54E-06 |
| **turquoise** | GO:0006091 | generation of precursor metabolites and energy | 0.10462 | 1.08E-35 |
| | GO:0051186 | cofactor metabolic process | 0.101902 | 1.31E-34 |
| | GO:0006631 | fatty acid metabolic process | 0.099185 | 2.13E-33 |
| | GO:0015980 | energy derivation by oxidation of organic compounds | 0.08288 | 4.35E-33 |
| | GO:0045333 | cellular respiration | 0.07337 | 2.40E-36 |
| **grey60** | GO:0002181 | cytoplasmic translation | 0.132075 | 2.43E-06 |
| | GO:0022613 | ribonucleoprotein complex biogenesis | 0.132075 | 0.008799 |
| | GO:0000028 | ribosomal small subunit assembly | 0.075472 | 9.07E-05 |
| | GO:0042274 | ribosomal small subunit biogenesis | 0.075472 | 0.00675 |
| | GO:0042255 | ribosome assembly | 0.075472 | 0.006768 |
| **magenta** | GO:0006397 | mRNA processing | 0.098684 | 0.000599 |
| | GO:0008380 | RNA splicing | 0.078947 | 0.003843 |
| | GO:0006641 | triglyceride metabolic process | 0.046053 | 0.003843 |
| | GO:0006639 | acylglycerol metabolic process | 0.046053 | 0.007832 |
| | GO:0006638 | neutral lipid metabolic process | 0.046053 | 0.007832 |
| **black** | GO:0006091 | generation of precursor metabolites and energy | 0.084444 | 1.46E-05 |
| | GO:0015980 | energy derivation by oxidation of organic compounds | 0.071111 | 1.36E-05 |
| | GO:0032868 | response to insulin | 0.057778 | 0.000206 |
| | GO:0005978 | glycogen biosynthetic process | 0.031111 | 0.000137 |
| | GO:0009250 | glucan biosynthetic process | 0.031111 | 0.000137 |
| **cyan** | GO:0007346 | regulation of mitotic cell cycle | 0.115942 | 0.036497 |
| | GO:0006520 | cellular amino acid metabolic process | 0.101449 | 0.012244 |
| | GO:0006631 | fatty acid biosynthesis process | 0.101449 | 0.036497 |
| | GO:1901605 | alpha-amino acid metabolic process | 0.086957 | 0.012244 |
| | GO:0043171 | peptide catabolic process | 0.043478 | 0.027279 |
| **brown** | GO:0051640 | organelle localization | 0.065455 | 0.033196 |
| | GO:0022613 | ribonucleoprotein complex biogenesis | 0.058182 | 0.033196 |

| | | | | |
|---|---|---|---|---|
| | GO:0034660 | ncRNA metabolic process | 0.054545 | 0.109875 |
| | GO:0042254 | ribosome biogenesis | 0.047273 | 0.042371 |
| | GO:0006338 | chromatin remodeling | 0.036364 | 0.033196 |
| **red** | GO:0022613 | ribonucleoprotein complex biogenesis | 0.097345 | 8.97E-08 |
| | GO:0042254 | ribosome biogenesis | 0.088496 | 1.68E-08 |
| | GO:0002181 | cytoplasmic translation | 0.070796 | 2.52E-12 |
| | GO:0042274 | ribosomal small subunit biogenesis | 0.044248 | 5.78E-07 |
| | GO:0042255 | ribosome assembly | 0.044248 | 9.70E-07 |
| **greenyellow** | GO:0006397 | mRNA processing | 0.114865 | 9.26E-06 |
| | GO:0008380 | RNA splicing | 0.081081 | 0.002137 |
| | GO:0022613 | ribonucleoprotein complex biogenesis | 0.081081 | 0.0045 |
| | GO:1903311 | regulation of mRNA metabolic process | 0.074324 | 0.000744 |
| | GO:0050684 | regulation of mRNA processing | 0.054054 | 0.002137 |
| **midnightblue** | GO:0006091 | generation of precursor metabolites and energy | 0.177419 | 4.20E-06 |
| | GO:0007005 | mitochondrion organization | 0.177419 | 8.22E-06 |
| | GO:0046034 | ATP metabolic process | 0.145161 | 6.11E-06 |
| | GO:0006119 | oxidative phosphorylation | 0.129032 | 2.72E-07 |
| | GO:0045333 | cellular respiration | 0.129032 | 5.95E-06 |
| **purple** | GO:0072594 | establishment of protein localization to organelle | 0.135135 | 1.56E-09 |
| | GO:0006457 | protein folding | 0.101351 | 1.22E-10 |
| | GO:0006278 | RNA-dependent DNA biosynthetic process | 0.060811 | 1.19E-07 |
| | GO:0007004 | telomere maintenance via telomerase | 0.060811 | 1.19E-07 |
| | GO:1904814 | regulation of protein localization to chromosome, telomeric region | 0.040541 | 1.19E-07 |
| **yellow** | GO:0034976 | response to endoplasmic reticulum stress | 0.132 | 1.49E-23 |
| | GO:0048193 | Golgi vesicle transport | 0.108 | 9.27E-16 |
| | GO:0006888 | endoplasmic reticulum to Golgi vesicle-mediated transport | 0.1 | 8.87E-22 |
| | GO:0035966 | response to topologically incorrect protein | 0.076 | 2.38E-13 |
| | GO:0006487 | protein N-linked glycosylation | 0.056 | 4.20E-12 |
| **tan** | GO:1903900 | regulation of viral life cycle | 0.060241 | 0.126771 |
| | GO:0050792 | regulation of viral process | 0.060241 | 0.126771 |
| | GO:0043903 | regulation of interspecies interactions between organisms | 0.060241 | 0.126771 |
| | GO:0044743 | protein transmembrane import into intracellular organelle | 0.036145 | 0.126771 |

| | | | | |
|---|---|---|---|---|
| | GO:0065002 | intracellular protein transmembrane transport | 0.036145 | 0.126771 |
| **lightcyan** | GO:0016125 | sterol metabolic process | 0.272727 | 5.68E-23 |
| | GO:1902652 | secondary alcohol metabolic process | 0.257576 | 3.01E-21 |
| | GO:0016126 | sterol biosynthetic process | 0.242424 | 2.86E-26 |
| | GO:1902653 | secondary alcohol biosynthetic process | 0.212121 | 6.39E-23 |
| | GO:0006695 | cholesterol biosynthetic process | 0.19697 | 5.00E-21 |
| **yellow** | GO:0048285 | organelle fission | 0.102857 | 1.69E-12 |
| | GO:0000280 | nuclear division | 0.094286 | 7.33E-12 |
| | GO:0044770 | cell cycle phase transition | 0.088571 | 4.38E-11 |
| | GO:0006260 | DNA replication | 0.071429 | 3.81E-11 |
| **yellow** | GO:0006778 | porphyrin-containing compound metabolic process | 0.037143 | 7.33E-12 |
| **turquoise** | GO:0006281 | DNA repair | 0.041486 | 5.43E-05 |
| | GO:0120031 | plasma membrane bounded cell projection assembly | 0.039628 | 0.001014 |
| | GO:0034660 | ncRNA metabolic process | 0.035913 | 0.001144 |
| | GO:0044782 | cilium organization | 0.029721 | 0.001014 |
| | GO:0019236 | response to pheromone | 0.014861 | 0.000473 |
| **brown** | GO:0070661 | leukocyte proliferation | 0.118785 | 8.50E-24 |
| | GO:0046651 | lymphocyte proliferation | 0.116022 | 8.50E-24 |
| | GO:0032943 | mononuclear cell proliferation | 0.116022 | 8.50E-24 |
| | GO:0030098 | lymphocyte differentiation | 0.11326 | 6.16E-19 |
| | GO:0070663 | regulation of leukocyte proliferation | 0.088398 | 7.90E-18 |
| **red** | GO:0042254 | ribosome biogenesis | 0.209877 | 1.55E-13 |
| | GO:0022613 | ribonucleoprotein complex biogenesis | 0.209877 | 2.09E-11 |
| | GO:0002181 | cytoplasmic translation | 0.17284 | 2.42E-16 |
| | GO:0042255 | ribosome assembly | 0.123457 | 2.09E-11 |
| | GO:0042274 | ribosomal small subunit biogenesis | 0.098765 | 2.15E-08 |
| **green** | GO:0006281 | DNA repair | 0.195876 | 9.73E-11 |
| | GO:0006260 | DNA replication | 0.134021 | 2.17E-08 |
| | GO:0009314 | response to radiation | 0.113402 | 0.000309 |
| | GO:0006302 | double-strand break repair | 0.103093 | 1.58E-05 |
| | GO:0010212 | response to ionizing radiation | 0.072165 | 0.000309 |
| **violet** | GO:0048285 | organelle fission | 0.102857 | 1.69E-12 |
| | GO:0000280 | nuclear division | 0.094286 | 7.33E-12 |
| | GO:0044770 | cell cycle phase transition | 0.088571 | 4.38E-11 |
| | GO:0006260 | DNA replication | 0.071429 | 3.81E-11 |
| | GO:0006778 | porphyrin-containing compound metabolic process | 0.037143 | 7.33E-12 |
| **royalblue** | GO:0006281 | DNA repair | 0.041486 | 5.43E-05 |

| | | | | |
|---|---|---|---|---|
| | GO:0120031 | plasma membrane bounded cell projection assembly | 0.039628 | 0.001014 |
| | GO:0034660 | ncRNA metabolic process | 0.035913 | 0.001144 |
| | GO:0044782 | cilium organization | 0.029721 | 0.001014 |
| | GO:0019236 | response to pheromone | 0.014861 | 0.000473 |
| **steelblue** | GO:0006397 | mRNA processing | 0.093711 | 7.86E-61 |
| | GO:0008380 | RNA splicing | 0.080764 | 1.14E-55 |
| | GO:0000375 | RNA splicing, via transesterification reactions | 0.059186 | 3.14E-42 |
| | GO:0000377 | RNA splicing, via transesterification reactions with bulged adenosine as nucleophile | 0.059186 | 3.14E-42 |
| | GO:0000398 | mRNA splicing, via spliceosome | 0.059186 | 3.14E-42 |
| **darkred** | GO:0042254 | ribosome biogenesis | 0.209877 | 1.55E-13 |
| | GO:0022613 | ribonucleoprotein complex biogenesis | 0.209877 | 2.09E-11 |
| | GO:0002181 | cytoplasmic translation | 0.17284 | 2.42E-16 |
| | GO:0042255 | ribosome assembly | 0.123457 | 2.09E-11 |
| | GO:0042274 | ribosomal small subunit biogenesis | 0.098765 | 2.15E-08 |
